# A neuroimaging database combining movie-watching, eye-tracking, sensorimotor mapping, and cognitive tasks

**DOI:** 10.1101/2025.09.25.678556

**Authors:** Egor Levchenko, Hugo Chow-Wing-Bom, Fred Dick, Adam Tierney, Jeremy I Skipper

## Abstract

We provide a multimodal naturalistic neuroimaging database (NNDb-3T+), designed to support the study of brain function under both naturalistic and controlled experimental conditions. The database includes high-quality 3T fMRI data from 40 participants acquired during full-length movie-watching and three sensory mapping tasks: somatotopy, retinotopy, and tonotopy. Each participant also completed synchronised eye-tracking during movie-watching and retinotopy, physiological recordings, and a battery of behavioural and cognitive assessments. Data were collected across two MRI sessions and a remote testing session, with all data organised in a BIDS-compliant format. Technical validation confirms high data quality, with minimal head motion, accurate eye-tracker calibration, and robust task-evoked activation patterns. The database provides a unique resource for investigating individual differences, functional topographies, multimodal integration, and naturalistic cognition. All raw and preprocessed data, quality metrics, and preprocessing scripts are publicly available to support reproducible research.

## Background & Summary

One of the main goals of human neuroscience is to uncover how the brain operates in the complex, continuous experiences during everyday life. To achieve this, researchers employ both naturalistic (e.g. watching a movie or listening to a narrative) and task-based paradigms (e.g. n-back task or sensory mapping tasks). Naturalistic paradigms tend to elicit higher immersion and attentiveness in participants (Ki et al., 2016), which improves ecological validity and also reduces head motion in the scanner (Vanderwal et al., 2019). The naturalistic approach has proven effective for studying a range of processes, including the hierarchy of temporal receptive fields (Lerner et al., 2011), event segmentation in memory (Baldassano et al., 2018), default mode network dynamics (Simony et al., 2016), selective attention (Nguyen et al., 2017), emotions and social cognition (Redcay & Moraczewski, 2020), and functional connectivity (Gal et al., 2022).

Over the past decade, numerous publicly available fMRI datasets have embraced naturalistic paradigms to explore how the brain responds to continuous, real-world stimuli. Notable examples include *StudyForrest* (Hanke et al., 2014), which combines 7T fMRI with eye-tracking and physiological recordings during audio and audiovisual presentation of the movie “Forrest Gump”; the Sherlock dataset (Chen et al., 2017), which includes free recall during scanning; and the *Grand Budapest Hotel* dataset (Visconti Di Oleggio Castello et al., 2020), focused on social cognition. Other large-scale efforts, such as the *Narratives* (Nastase et al., 2021) and *CamCAN* (Shafto et al., 2014) datasets, have used spoken stories or short films to investigate ageing, language, and attention. These resources have advanced the field by demonstrating that naturalistic stimuli evoke reliable, temporally aligned neural responses across individuals, and can be used to study phenomena like event segmentation, narrative comprehension, and social perception. Despite this progress, most existing datasets emphasise either naturalistic stimulation or controlled task-based mapping, rarely integrating both within the same cohort. Furthermore, multimodal recordings such as eye-tracking and physiological measures are often missing or only partially available.

Among the currently available naturalistic fMRI datasets, the *Naturalistic Neuroimaging Database* (NNDb v1.0) (Aliko et al., 2020) is notable for a large sample (N=86) and extensive behavioural and cognitive phenotyping for each participant who watched one of ten full-length feature films spanning diverse genres. While NNDb v1.0 was designed to emphasise diversity of movie genres, the present dataset, the *Naturalistic Neuroimaging Database 3T+* (NNDb-3T+), provides a uniquely rich multimodal resource with several tasks and physiological recordings for each participant. The dataset combines whole-brain fMRI 3T data (2×2×2mm resolution) from 40 participants during full-length movie-watching (*Back to the Future)*, somatotopy, retinotopy, and tonotopy tasks, eye-tracking data, physiological recordings, and extensive behavioural measures. Figure 1 provides an overview of the collected data, preprocessing techniques, and quality control analyses for each task. NNDb-3T+ is, to our knowledge, the first publicly available dataset to combine naturalistic movie viewing, three distinct sensory mapping tasks, synchronised eye-tracking, physiological monitoring, and behavioural profiling within the same participants. We anticipate that NNDb-3T+ will support a wide range of future investigations, including individual differences in naturalistic processing, neural modelling of sensory hierarchies, and the development of novel analytic methods for multimodal neuroimaging. All raw and preprocessed data are shared in Brain Imaging Data Structure (BIDS) valid format (Gorgolewski et al., 2016), and quality control metrics are provided. Scripts for preprocessing and validation are openly available on GitHub to promote reproducibility (https://github.com/levchenkoegor/movieproject2).

**Figure 1.**
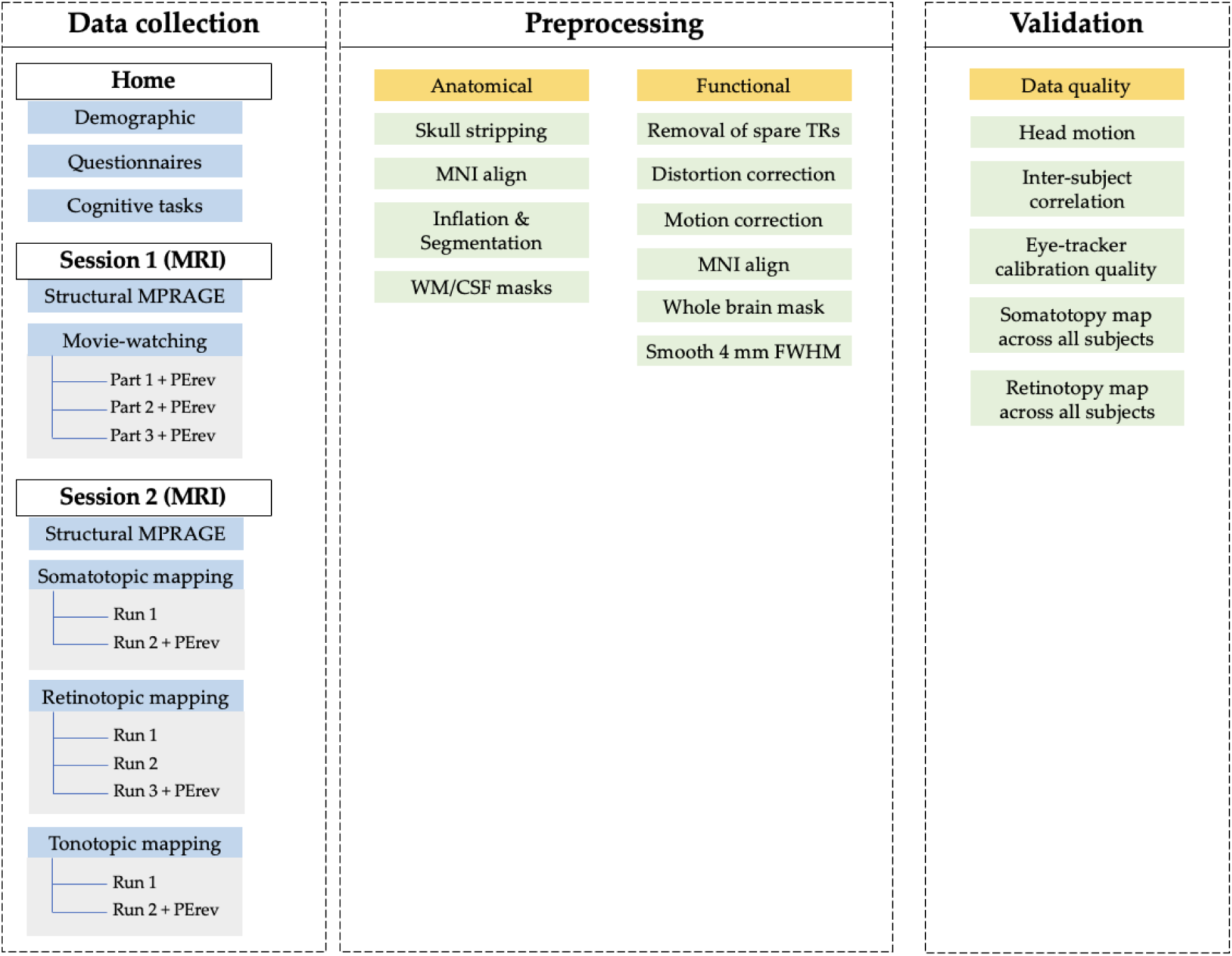
Overview of data collection, preprocessing, and validation pipeline. (1) Data collection: data were acquired across two MRI sessions and a home assessment. At home, participants completed demographic questionnaires, cognitive tasks, and self-report measures. Session 1 included acquisition of a 5-minute high-resolution structural MPRAGE scan and a naturalistic movie-watching task split into three parts, each accompanied by a phase-encoding reversed (PErev) scan for distortion correction. Session 2 comprised 2-minute structural imaging and functional localizers: somatotopic mapping (2 runs, one with PErev), retinotopic mapping (3 runs, including PErev), and tonotopic mapping (2 runs, including PErev). (2) Preprocessing: the pipeline was applied to each task separately. Somatotopy and movie-watching (Back to the Future) tasks followed the same pipeline shown in the current Figure, and retinotopy and tonotopy tasks followed minimal preprocessing (no alignment, masking and smoothing; see Preprocessing for more details). (3) Validation: Data quality and validity were assessed using multiple metrics: amount of head movement for each task, inter-subject correlation during the movie task, eye-tracker calibration quality before each run of the movie, and averaged activation maps for somatotopy and retinotopy tasks.

## Methods

### Participants

We recruited 44 participants using participant pool management software (http://www.sona-systems.com/), advertisement on the University campus, and word of mouth. Participants were prescreened based on MRI safety criteria (e.g., no metal implants) and self-reported good (or corrected to good) vision and hearing. The final sample consisted predominantly of right-handed, native English speakers, between 18 and 45 years old with no neurological diseases. A few exceptions are noted: one left-handed person with German/English first language and one with Farsi, one ambidextrous, five participants reported psychological or psychiatric diagnoses (Obsessive Compulsive Disorder, Autism Spectrum Disorder, Generalised Anxiety Disorder, Depression and ADHD). Two participants failed to attend and two were excluded due to technical problems during the acquisition (e.g., felt uncomfortable inside the scanner). The final sample consisted of 40 participants: 21 females, 18–45 years, M = 29.02, SD = 6.20 years (though one participant did not complete the questionnaire).

The questionnaire and cognitive tasks collected remotely were approved by the Ethics Committee of the School of Psychological Sciences at Birkbeck (Reference number: 2324006). The whole study including MRI was approved by the ethics committee at University College London (Reference number: fMRI/2023/003). All participants provided written consent to participate in the study and share their data. At the end of the study, participants received £67.50 in the form of a voucher.

### Procedure

Before their visit, participants filled out an MRI safety form. If they were MRI-safe, they filled out a questionnaire about demographic information, language background, musical experience, and knowledge of movies. Then the participant completed a set of cognitive tests on the *Cognitron* platform (https://www.cognitron.co.uk/) and two scanning sessions were scheduled. Two consent forms were signed online: one before the questionnaire and one before the cognitive tests.

At the beginning of the first scan day, the participant completed an MRI safety form again, and signed a paper-based ethics consent form. The participant was screened by an MRI operator once more before entering the scanner room. If the participant was deemed completely safe to go inside the scanner, we initiated *Session 1*.

Once in the scanning room, the participant chose suitable earbud sizes for noise-attenuating headphones and donned a hairnet. The participant then put the earbuds in and was helped to lie back into a novel head motion reduction device (’MR-MinMo’, patent number GB 2205139.5 filed on 07 April 2022). The device consists of a frame that sits within the head coil with a set of inflatable and passive cushions to reduce head motion and increase participant comfort. An additional knee pillow was put under the participant’s legs for comfort and to minimise body movements.

Once the participant was moved inside the scanner bore, the in-bore light was turned off, as were the overhead lights in the scanner room. Next, a short segment of the movie was played to ensure the participant could see the video and hear the audio clearly. Audio volume was adjusted for each participant separately. A localizer was then run, and the slab was adjusted to capture the whole brain. If it was not possible then the operator prioritised removing as few slices of the cerebellum as possible. When ready, the presentation script was started and eye-tracker calibration and validation procedures were completed. The operator made adjustments to achieve the best validation quality for the eye-tracker, and then proceeded to the movie-watching task.

During *Session 1*, the participant watched the entirety of ‘Back To The Future’ (*backtothefuture)*, (Zemeckis, 1985), divided into three parts (see below for timing). Eye-tracker calibration was performed before each part. The entire process for *Session 1* took about three hours.

In *Session 2*, we followed the same procedures for participant preparation. Participants completed several different tasks inside the scanner during *Session 2*: 1) *somatotopic mapping*, where the participant performed prescribed movements inside the scanner; 2) *retinotopic mapping*, where different checkerboard patterns were presented while the participant was instructed to fixate on the dot in the middle of the screen, responding every time the dot changed colour; and 3) *tonotopic mapping*, during which a series of tone sequences were played, with participants to press a button when heard a repeated sequence (see ‘*Tasks’* for further details). The entire process for *Session 2* took about two hours.

### Tasks

Some participants failed to come back for the second day of testing and some tasks were not included in the final dataset due to technical reasons. The number of tasks completed by each participant is shown in *Table 1* below.

**Table 1.** The number of tasks completed by each participant. The ‘+’ means that the participant fully completed the task, and the ‘-’ means that the data is missing or the participant didn’t complete the task.

| sub id | Questionnaires | Cognitive tasks | Movie-watching | Somatotopy | Retinotopy | Tonotopy |
| --- | --- | --- | --- | --- | --- | --- |
| sub-01 | + | + | + | + | + | + |
| sub-02 | + | + | + | + | + | + |
| sub-03 | + | + | + | + | + | + |
| sub-04 | + | - | + | - | - | - |
| sub-05 | + | + | + | + | + | - |
| sub-06 | + | + | + | + | + | - |
| sub-07 | + | - | + | + | + | + |
| sub-08 | - | - | + | - | - | - |
| sub-09 | + | + | + | + | + | + |
| sub-10 | + | + | + | + | + | + |
| sub-11 | + | - | + | + | + | + |
| sub-12 | + | + | + | + | + | + |
| sub-13 | + | - | + | + | + | + |
| sub-14 | + | + | + | + | + | + |
| sub-16 | + | + | + | + | + | + |
| sub-17 | + | + | + | + | + | + |
| sub-18 | + | + | + | + | + | + |
| sub-19 | + | + | + | + | + | + |
| sub-20 | + | + | + | + | + | + |
| sub-21 | + | + | + | + | + | + |
| sub-22 | + | + | + | + | + | + |
| sub-23 | + | + | + | + | + | + |
| sub-24 | + | + | + | + | + | + |
| sub-25 | + | + | + | + | + | + |
| sub-26 | + | + | + | + | + | + |
| sub-27 | + | + | + | + | + | + |
| sub-29 | + | + | + | + | + | + |
| sub-30 | + | + | + | + | + | + |
| sub-31 | + | + | + | + | + | + |
| sub-32 | + | + | + | + | + | + |
| sub-33 | + | + | + | + | + | + |
| sub-35 | + | + | + | + | + | + |
| sub-36 | + | + | + | + | + | + |
| sub-37 | + | + | + | + | + | + |
| sub-38 | + | + | + | + | + | + |
| sub-39 | + | + | + | + | + | + |
| sub-40 | + | + | + | + | + | + |
| sub-42 | + | + | + | + | + | + |
| sub-43 | + | + | + | + | + | + |
| sub-44 | + | + | + | + | + | + |
| **Total N** | **39** | **35** | **40** | **38** | **38** | **36** |

### Home

#### Questionnaires

The entire questionnaire took approximately 30 minutes to complete. The first section consists of questions regarding basic demographics, language proficiency, and background in music and movie preferences. The second section included 9 validated psychological questionnaires. The set of questionnaires was selected to comprehensively assess participants’ mental health, well-being, inner experience, self-talk, mindfulness, and awareness. The selected measures are widely validated, reliable, and efficient, ensuring that they capture a broad spectrum of psychological functioning. These were implemented on the Qualtrics platform (https://qualtrics.ucl.ac.uk), and are described next.

The *Patient Health Questionnaire (PHQ)* is a 9-item tool for diagnosing depression and various other mental health conditions frequently seen in primary care settings (Kroenke et al., 2001). Each item has a scale from 0 (not at all) to 3 (nearly every day). It is notably shorter than many other depression assessments, yet it maintains similar levels of sensitivity and specificity.

The 7-item scale for *General Anxiety Disorder (GAD-7)* is a valid and efficient instrument to screen and evaluate the severity of the condition in both clinical and research settings (Spitzer et al., 2006). Each item has a scale from 0 (not at all) to 3 (nearly every day). GAD is among the most frequently observed anxiety disorders in both general medical practice and the broader population.

The *Warwick Edinburgh Mental Well-Being Scale (WEMWBS)* is a widely used measure of mental well-being, comprising exclusively positively worded items (Tennant et al., 2007). Participants evaluate their mental well-being over the last two weeks. The scale has 14 questions with a 5-point Likert scale (none of the time, rarely, some of the time, often, all of the time).

The *Nevada Inner Experience Questionnaire (NIEQ)* is used to assess the subjective frequency of experiencing five common phenomenological categories of inner thought (inner speaking, inner seeing, unsymbolized thinking, feelings, and sensory awareness) via a visual analogue scale (Heavey et al., 2019). The NIEQ has 10 items with two types of questions: ‘How frequently…?’ with a scale from 0 (never) to 100 (always) and ‘Generally speaking, what portion…?’ with a scale from 0 (none) to 100 (all).

The *Self Talk Scale (STS)* is used to assess the subjective frequency at which people engage in various modes of self-talk (Brinthaupt et al., 2009). The modes of self-talk assessed by this scale, as delineated by a four-factor structure, relate to ‘self-regulatory’ elements, including social assessments, self-criticism, self-reinforcement and self-management. Each question starts with ‘I talk to myself when…’ and the participant needs to evaluate the frequency on a 5-point scale (1 - never, 5 - very often).

The *Varieties of Inner Speech Questionnaire - Revised (VISQ-R)* is used to assess participants’ subjective frequency and phenomenological characteristics of their experience of inner speech (Alderson-Day et al., 2018). The characteristics comprise a four-factor model, with factors comprising: dialogical inner speech, condensed inner speech (as compared to spoken aloud), experience of other people’s voices, and self-evaluative inner speech. It consists of 26 items and each item is rated on a scale of 1 (never) to 7 (all the time).

The *Five Facets of Mindfulness Questionnaire (FFMQ)* is a 39-item instrument that uses five polytomous response options to assess five different aspects of mindfulness: observing, describing, acting with awareness, non-judging, and non-reactivity to inner experience (Baer et al., 2008). Each item is rated on a scale of 1 (Never or very rarely true) to 5 (Very often or always true).

The *Multidimensional Assessment of Interoceptive Awareness version II (MAIA-II)* evaluates eight factors of interoceptive body awareness: noticing, not-distracting, not-worrying, attention regulation, emotional awareness, self-regulation, body listening and trusting (Mehling et al., 2018). It consists of 37 items and each item is rated on a scale of 0 (never) to 5 (always).

The *White Bear Suppression Inventory (WBSI)* is a 15-item questionnaire measuring thought suppression (Wegner & Zanakos, 1994). Chronic thought suppression is a variable that is related to obsessive thinking and negative affect associated with depression and anxiety. Each item is rated on a 5-point scale from strongly disagree (1) to strongly agree (5).

#### Cognitive tasks

The battery of cognitive tasks was designed to comprehensively assess a range of cognitive abilities, including memory, executive function, attention, reasoning, and creativity. We selected tasks to sample key domains of cognition relevant to general intelligence, cognitive flexibility, and problem-solving, providing a robust framework for evaluating individual differences in cognitive function. We used 16 different tasks from the *Cognitron* platform. A detailed description of each task is available in the original publication and supplementary material by the *Cognitron* team (Del Giovane et al., 2023). The participants completed all the tasks remotely. The whole battery took around 45 minutes to complete. A description of each task is provided next in the order presented to the participant. Figures illustrating the trial structure of each task are available in the *Supplementary Materials*.

##### Object Memory Immediate and Delayed

This task measures memory capacity on short and long-term scales. Participants viewed a list of 20 black-and-white objects (for example, stairs, table, ladle, etc.), presented once at a time. They were instructed to remember as many as they could. In the *Immediate* version of the task, participants’ short-term memory was assessed by presenting a grid containing one object from the previously presented set along with 7 similar “distractor” objects. Participants had to identify and click the object they recognised from the original set. In total, 20 grids were presented (see Supplementary Fig. 1). Participants repeated this task at the end of the battery of cognitive tasks to assess long-term memory (*Delayed* version).

##### Word Memory Immediate and Delayed

This task is similar to the *Object Memory Immediate and Delayed* but uses words instead of images (see Supplementary Fig. 11).

##### 2D manipulations

This task measures the ability to mentally rotate a grid within a two-dimensional space. Participants were presented with a target grid partially filled with coloured squares, as well as four comparison grids. One of these grids was a rotated transformation of the target grid (see Supplementary Fig. 2). Participants were instructed to identify the rotated grid as quickly and accurately as possible.

##### Intra/extra-dimensional set-shifting task (ID/ED)

The ID/ED task is a computerised analogue of the Wisconsin Card Sorting Task (Grant & Berg, 1948), designed to assess cognitive flexibility. Participants were presented with four squares, followed by two objects appearing in two randomly selected squares. They were instructed to identify the underlying rule by clicking on the object that matched the current rule; the rule changed after a number of correct responses (see Supplementary Fig. 3). There were two main types of rule changes: (1) an ID rule where the rule continues to rely on the same dimension (e.g., shape), but the the shape changes (e.g., from triangle vs. circle to square vs. star) and (2) an ED rule where the rule changes to a different dimension altogether (e.g., from selecting based on shape to selecting based on line pattern). These shifts require increasing levels of cognitive flexibility, with ED shifts being particularly challenging because they demand a shift of attention to an entirely new dimension. Performance on ED trials is therefore considered a strong indicator of flexible thinking and attentional control.

##### Spatial Span

This task assesses visuospatial working memory capacity. Participants were presented with a 4-by-4 grid in which a sequence of squares lit up and were asked to repeat the presented sequence by clicking on the squares in the order they lit up. The sequence began with the two squares, and after each correct response, the sequence length increased by one square (see Supplementary Fig. 4).

##### Digit Span

The task is designed to assess verbal working memory. Participants were presented with a sequence of digits and asked to recall them by clicking on the digits in the correct order (see Supplementary Fig. 5). When a sequence was reproduced correctly, the next sequence length was increased by one digit.

##### Switching Stroop

This task is a modified version of the classical Stroop test (Stroop, 1935) and incorporates a switching condition in addition to the classic interference condition. On each trial, participants were presented with the cue words ‘Text’ or ‘Ink’ accompanied by two coloured words, ‘RED’ and ‘BLUE’ and a central coloured box - the ‘Ink’ (red or blue). Based on the given instruction (“Text” or “Ink”), participants were required to click on one of the two words. If the rule was ‘Ink’, participants selected the word printed in the same ink colour as the central box. If the rule was ‘Text’, they selected the word whose meaning matched the colour of the central box (see Supplementary Fig. 6).

##### Verbal Reasoning

This task measures participants’ ability to interpret and analyse written material. Participants read syntactically more complex sentences conveying different thematic relations. In each trial, a picture of a square and a circle was shown along with the written sentence (for example, ‘the square is contained by a circle’). Participants responded true or false via mouse (see Supplementary Fig. 7).

##### Beads task

This task measures impulsivity. Participants were instructed to work out which of two bead colours is the most prevalent in the jar. Participants could reveal one bead at a time by clicking the button labelled ‘Reveal a bead’. At any point, they could choose to guess the dominant colour in the jar by clicking on one of the colours (see Supplementary Fig. 8).

##### Alternative Use Task

This task measures creativity. Participants were asked to invent as many alternative uses for a common item as possible (for example, a newspaper). After each response, participants were asked if the idea came to their mind as an ‘Aha’ moment. The task was limited to two minutes or 20 alternatives.

##### Divergent Association task

This task measures creativity and specifically divergent thinking. Participants were asked to think of 10 words in four minutes that were as different from each other as possible.

##### Verbal analogies

This task assesses verbal reasoning and the ability to understand and apply logical relationships between word pairs. Participants were presented with a statement where the relationship between two pairs of words must be assessed as either ‘true’ or ‘false’. It followed the structure: ‘A is to B as C is to D’. Participants must determine whether the relationship between A and B was indeed analogous to the relationship between C and D (see Supplementary Fig. 9).

##### Word Definitions

This task estimates the vocabulary size and language comprehension level. Participants needed to choose the correct definition of the word out of four options (see Supplementary Fig. 10).

##### Spotter (digit vigilance)

This task assesses the participant’s vigilance. Participants briefly observed a sequence of numbers, obscured by ‘noisy’ pixels. The task required the participant to spot and click anywhere on the screen as soon as they recognised a zero (‘0’). Occasionally, the participant was asked how motivated and tired they felt on a scale from one (not at all) to six (extremely).

### Session 1

Participants watched the movie ‘Back To The Future’ (Zemeckis, 1985). The duration of the movie was one hour 51 minutes and 14 seconds. They were told to remain still inside the scanner and enjoy watching the movie.

The movie file was cropped into three runs using the ‘ffmpeg’ package:

*ffmpeg -ss 00:00:00 -i back_to_the_future.mp4 -c copy -t 00:33:48 back_to_the_future_cut1-34min.mp4*
*ffmpeg -ss 00:33:36 -i back_to_the_future.mp4 -c copy -t 00:37:39 back_to_the_future_cut2-38min.mp4*
*ffmpeg -ss 01:11:03 -i back_to_the_future.mp4 -c copy -t 01:00:00 back_to_the_future_cut3-40min.mp4*

The specific time for each cut was chosen to maintain a similar duration between runs and to retain a smooth transition between scenes. The 2nd and the 3rd runs included 12 seconds of the scene from the previous run to provide enough time for the hemodynamic response function (HRF) and psychological functioning to (theoretically) recover to a state similar to that in the preceding run. Each run started with eight dummy TRs. Thus, eight, 16 and 16 TRs were removed during the analysis of runs 1-3, respectively such that the fMRI data matched the length of the full movie, without overlaps. The length of the resulting runs was 34 minutes, 38 minutes three seconds and 40 minutes 12 seconds for runs one, two, and three respectively (1360, 1522 and 1608 TRs).

The cropped files maintained the original video size and quality, using all frames with no cropping or other transformations:

- Video (codec): H.264 (High)
- Audio (codec, sampling rate, bitrate, channels): AAC (LC), 48.0 kHz, 339 kbps, 5.1
- Resolution (pixels): 720 x 576
- Aspect Ratio: 16:9
- Frame rate (fps): 25

The movie presentation was implemented using a script executed from MATLAB (9.13.0.204977 R2022b) using PsychToolBox (v. 3.0.18) on a Windows PC (Windows 11 Pro v22H2, 64-bit operating system) with GStreamer (v. 1.0).

### Session 2

#### Somatotopic mapping

Each participant underwent a training session before going inside the scanner. The participant watched short videos depicting each movement, listened to the instructions from the researcher and practised to perform the movement. The researcher assessed the movement and if all of them were performed well, the participant began the scanning session. Participants did one more short training session inside the scanner to find the most comfortable limb positions to perform the movements properly. The researcher assessed the quality of movements through the in-bore camera. When training was completed, the participant underwent two runs of the block design experiment.

Each of the 8 experimental conditions involved moving one of four body parts (hand, foot, face, and tongue) on the left or right side. The pattern for each movement is outlined in Table 2 below.

**Table 2.** Somatotopic mapping task. The experimental conditions and pattern for each movement.

| Experimental condition (movement) | The pattern of the movement |
| --- | --- |
| Left/right hand | Clench and relax the fist |
| Left/right foot | Flex and extend the toes |
| Left/right part of the face | Pull the corner of the mouth down and to the side |
| Left/right tongue | Touch the last molar with the tip of the tongue when the mouth and jaw are closed |
| Rest | No movement, remain still |

In each condition, the participant maintained eye fixation on the screen where the instructions were presented. The instructions consisted of one line of text with a specific movement (for example, ‘right foot’). The participant kept doing the movement until instructions were changed on the screen.

During the run, a metronome was played at 30bpm. The participants were instructed to pace the movement to the metronome clicks.

The sequence of runs was counterbalanced across participants. Runs one and two lasted for seven minutes 47 seconds and seven minutes 49 seconds respectively (excluding eight spare TRs in the beginning). The duration of each movement varied between 15 and 22 seconds and was repeated three times per run. The sequence of movements was pseudorandomised (exact sequences for each run are shown in the *Supplementary Materials*). The experiment was implemented in PsychoPy (v. 2023.2.3) using standard builder functionality.

#### Retinotopic mapping

To map how the visual space is systematically represented across cortical areas (i.e., retinotopic organisation), we used a population receptive field (pRF) mapping task during fMRI scanning (Dumoulin & Wandell, 2008). The stimulus consisted of a black-and-white contrast-reversing checkerboard (2Hz), embedded within a rotating wedge (20° angle) and expanding/contracting ring. Each run included six ring cycles (48 seconds each, logarithmic eccentricity scaling) and eight wedge cycles (36 seconds each, alternating clockwise/anticlockwise). Stimuli covered a maximum eccentricity of 8.6 from fixation and updated position every one second/TR.

Baseline periods (20 seconds) were inserted at the start, midpoint, and end, during which participants fixated on a central dot against a mid-grey background. A small white fixation dot (0.2° visual angle radius) and a black radial grid were always visible to support stable fixation. Each run lasted 348 seconds (five minutes 56 seconds), and participants completed three identical runs.

To maintain engagement, participants performed a simple detection task, pressing a button whenever the fixation dot changed from white to black. Due to technical issues, behavioural responses from the first nine participants were not recorded. Eye movements were tracked using an Eyelink 1000 Plus (SR Research, Ottawa, ON), with a 5-point custom calibration performed before each run. The task was implemented in MATLAB using PsychToolBox (v. 3.0.18) on a Windows PC (Windows 11 Pro v22H2, 64-bit operating system) with GStreamer (v 1.0).

#### Tonotopic mapping

The full description of the task can be found in the original study on tonotopic mapping of the auditory cortex (Dick et al., 2017). Participants listened to four-tone motifs and performed a one-back task on infrequent repeats. The full frequency range of the tones (175-5286 Hz) was divided into ten spectrally delimited bands, with each band having a 6-semitone range. At any given point in time, tones were selected from only one frequency band; after 10 motifs, the frequency range stepped up (in one run) or down (in the other) to the next band. Each run swept through the full frequency range four times (64 seconds per sweep). In this way, each frequency band occurred with consistent timing within a sweep; critically, then, voxels that respond preferentially to that frequency range should also respond at a consistent phase lag (Dick et al., 2012; Dick et al., 2017; Sereno et al., 1995).

The participants were instructed to press the button every time they heard the same tone twice in a row (1-back task). The experiment was implemented in PsychoPy (v. 2023.2.3) using standard builder functionality.

### Data acquisition

For both sessions and all tasks, functional and anatomical images were acquired on a 3.0T Siemens MAGNETOM Prisma using a custom-modified 30-channel radio-frequency (RF) head coil (Siemens Healthcare, Erlangen, Germany). To facilitate wide-angle viewing, this version of the head coil has had the two eye coil elements and associated housing removed, but is otherwise identical to the typical 32-channel coil.

#### MRI parameters: backtothefuture task

We used the CMRR (Xu et al., 2013) multiband echo-planar imaging sequence with TR = 1500 ms, TE = 35.2 ms, 72 interleaved slices, slice thickness 2.0 mm, voxel size 2 mm isotropic, anterior-posterior phase encoding direction (A >> P), field of view 212 mm, flip angle 60 deg, echo spacing 0.56ms, bandwidth 2620 Hz/Px, with a 4x multiband acceleration factor. The first, second, and third run had 1360, 1522, and 1608 TRs respectively (including eight spare TRs at the beginning of each run). A phase reverse encoding (P >> A) scan was acquired right after each run.

A 5-min high-resolution T1-weighted MPRAGE anatomical MRI scan followed the functional scans (TR = 2300 ms, TE = 2.98 ms, 208 sagittal slices, slice thickness 1.0 mm, voxel size 1 mm isotropic, anterior-posterior phase encoding direction (A >> P), field of view 212 mm, flip angle 9 deg, echo spacing 7.1 ms, bandwidth 240 Hz/Px).

#### MRI parameters: somatotopic mapping

At the beginning of *Session 2*, a structural scan was acquired to help position the field of view more accurately for upcoming tasks. A 2-min high-resolution T1-weighted MPRAGE anatomical MRI scan was acquired (TR = 1530 ms, TE = 2.98 ms, 176 sagittal slices, slice thickness 1.0 mm, voxel size 1.0 mm isotropic, anterior-posterior phase encoding direction (A >> P), field of view 256 mm, flip angle 9 deg, echo spacing 7.1 ms, bandwidth 240 Hz/Px, GRAPPA=4).

We used multiband echo-planar imaging sequence (TR = 1000 ms, TE = 30.0 ms, 44 interleaved slices, slice thickness 2.0 mm, voxel size 2 mm isotropic, anterior-posterior phase encoding direction (A >> P), field of view 212 mm, flip angle 62 deg, echo spacing 0.7 ms, bandwidth 1814 Hz/Px) with a 4x multiband acceleration factor (Xu et al., 2013). The first and the second run had 467 and 469 TRs respectively (including eight spare TRs at the beginning of each run). A phase reverse encoding (P >> A) scan was acquired at the end of the task.

#### MRI parameters: retinotopic mapping

We used multiband echo-planar imaging (TR = 1000 ms, TE = 35.2 ms, 48 interleaved slices, slice thickness 2.0 mm, voxel size 2 mm isotropic, anterior-posterior phase encoding direction (A >> P), field of view 212 mm, flip angle 60 deg, echo spacing 0.56 ms, bandwidth 2620 Hz/Px) with a 4x multiband acceleration factor. All three runs had the same number of 356 TRs (including eight spare TRs at the beginning of each run). A phase reverse encoding (P >> A) scan was acquired at the end of the task.

#### MRI parameters: tonotopic mapping

We used multiband echo-planar imaging with the same parameters as for the *somatotopic mapping* task (see above). Both runs had the same number of 264 TRs (including eight spare TRs at the beginning of each run). A phase reverse encoding (P >> A) scan was acquired at the end of the task.

##### Eye-tracker apparatus

Eye tracking data were acquired using an MRI-compatible EyeLink 1000 Plus long-range mount (SR Research Ltd., Mississauga, Ontario, Canada). The optic camera head (f=50mm/F1.4 lens) and illuminator (FL-890) were positioned horizontally behind the screen inside the bore on a tray. A front silvered mirror was placed on the anterior MRI coil. The binocular setup was used with a sampling frequency of 1000 Hz and a 9-dot calibration procedure covering the whole screen (5-dot calibration for the *retinotopy* task). For accuracy validation, participants had to fixate on the same dots as during calibration (EyeLink software v. 5.15). The calibration and validation procedures were repeated until the best possible accuracy was achieved.

The setup inside the bore was the same between days. Stimuli were presented in full-screen mode through a mirror-reversing LCD projector (EPSON LB-1100U) to a rear-projection screen, with participants viewing through the front silvered mirror attached to the head coil. Participants were positioned 57.5 cm from the screen, which was viewed via a mirror attached to the head coil, and measured 35.5 cm in width and 26 cm in height. Stimuli were presented in their native resolution and subtended 28.9° × 18.3° (29.6×18.5cm) of visual angle. The eye-tracker camera and illuminator were located inside the bore behind the screen on a movable platform. The position of the screen, projector, eye-tracker platform, and first-surface mirror was controlled between participants and checked to be the same.

The eye tracker data were acquired during *backtothefuture* and *retinotopy* tasks. The MATLAB script started with the calibration and validation of the eye-tracker, and if the calibration was deemed sufficient, the main experiment began after 8 spare TRs were received from the scanner. The script sent messages (‘MSG’ in ASCII eye link data file) indicating the beginning of the event. In the *backtothefuture* task, at the beginning of the movie, each pulse and frame was sent to the ASCII data file (see *Data Records* for more details). In both tasks, the presentation script was implemented in MATLAB (9.13.0.204977 R2022b) using PsychToolBox (v. 3.0.18) on a Windows PC (Windows 11 Pro v22H2, 64-bit operating system) with GStreamer (v 1.0).

##### Physiological recordings

The pulse oximetry data were acquired using a Siemens wireless peripheral pulse unit (PPU; Siemens Healthcare GmbH, Erlangen, Germany). The PPU sensor was attached to the index finger of the left hand. Participants were instructed to avoid any left-hand movements (including fingers) to prevent movement artefacts. The pulse data was acquired during *backtothefuture*, *retinotopy* and *tonotopy* tasks.

##### Preprocessing

The raw DICOM data were transformed to a Brain Imaging Data Structure (BIDS) valid format using the *heudiconv* tool (https://github.com/nipy/heudiconv; v. 1.3.0) (Gorgolewski et al., 2016). All anatomical images were defaced using the *pydeface* tool (https://github.com/poldracklab/pydeface; v. 2.0.2). MRI data were preprocessed using the AFNI software suite (AFNI_23.0.03 ‘Commodus’) to ensure data quality and prepare it for statistical analysis (Cox, 1996; Cox & Hyde, 1997).

The *backtothefuture* and *somatotopy* tasks were analysed using the same preprocessing pipeline and the *tonotopy* and *retinotopy* tasks were preprocessed differently according to the pipeline described in the previous relevant papers (Chow-Wing-Bom et al., 2025; Dekker et al., 2019; Dick et al., 2017).

##### Anatomical

The anatomical T1-weighted image was skull-stripped and then nonlinearly aligned to the MNI152 2009 template, which generated a standard-space anatomical image, a nonlinear warp and an affine transformation matrix (using AFNI *SSwarper* command). The anatomical surfaces were reconstructed using Freesurfer software (*recon-all* with default parameters, version 7.3.2-20220804-6354275, http://www.freesurfer.net) (Destrieux et al., 2010; Fischl, 2012). The resulting surfaces were converted to AFNI friendly format (using Surface Mapper (SUMA) tool) and later used to create white matter and ventricle regions of interest to use them as nuisance regressors during preprocessing. These regions were eroded and used as noise regressors in the preprocessing of *backtothefuture* and *somatotopy* tasks.

##### Functional

Functional data underwent multiple preprocessing steps to ensure alignment and reduce artefacts. First, the initial volumes of each functional run were removed to eliminate pre-steady-state effects. The first eight TRs were removed from run 1, and the first sixteen TRs were removed from runs 2 and 3 (*backtothefuture* task). In other tasks, the first eight TRs were removed from each run. To correct for geometric distortions caused by susceptibility-induced field inhomogeneities, a blip-up/blip-down correction was applied using a reverse-phase encoding field map. The median images of the forward and reverse phase-encoding runs were extracted, and their midpoint warps were computed and applied to the functional runs using *‘3dNwarpApply*’.

Motion correction was performed using a two-pass alignment strategy (‘*3dvolreg*’). First, volumes within each run were aligned to the run-specific reference; then, these within-run bases were themselves aligned to a common reference volume. This hierarchical strategy allowed all runs to be brought into a shared alignment space. To further improve robustness, alignment was performed using a two-pass procedure: an initial low-resolution stage estimated gross head motion, which was then refined at full resolution. Those procedures were applied to all tasks, and all further steps described below were applied only to *backtothefuture* and *somatotopy* tasks, but not to the *tonotopy* task. The specifics of processing for the *retinotopy* task are described in the section below (Retinotopic mapping).

Each volume was aligned to a reference volume determined by the minimum outlier fraction across all runs. The aligned functional images were then aligned to the anatomical image (‘*align_epi_anat.py*’). Finally, the functional data were nonlinearly aligned to MNI standard space.

A whole-brain mask was generated by computing the union of individual run-based masks. The functional images were then smoothed using a 4 mm full-width half-maximum (FWHM) Gaussian kernel to improve signal-to-noise ratio (‘*3dBlurToFWHM*’). Next, the data were temporally band-pass filtered (0.01-1 Hz) to reduce low-frequency drifts and high-frequency noise.

Because the runs of the *backtothefuture* task were very long, the baseline polynomial degree was fixed to two. For all other tasks, the degree was computed automatically using AFNI algorithms based on the length of the run.

##### Retinotopic mapping

To enable surface-based analysis, each participant’s functional data were first aligned to their high-resolution anatomical image using the following approach. As described previously, the functional volumes used for retinotopic mapping were motion-corrected using a two-pass alignment strategy (3dvolreg), which included within-run and across-run registration to a common reference volume. Importantly, this motion correction was performed before any additional preprocessing (e.g., spatial blurring or nuisance regression), so the volumes used here reflect unblurred, motion-corrected data.

Each participant’s motion-corrected volume was registered to their high-resolution anatomical image using boundary-based registration (bbregister, FreeSurfer). Registration quality was visually inspected and quantified with the cost function value. All participants with a minimum cost function value above 0.45^1^ were re-registered to a single-band reference volume from run one. This applied to participants 01, 07, 20, 21, 25 and 44.

Following successful registration, volumetric data for each run were then resampled onto the surface of both hemispheres using cortical surface models reconstructed from each participant’s high-resolution anatomical scan (*mri_vol2surf* command from Freesurfer). Sampling was performed at 50% cortical depth, using a 3 mm FWHM surface-based smoothing kernel to enhance signal-to-noise ratio while preserving spatial specificity. This step generated a surface-based time series for each hemisphere and run, enabling subsequent retinotopic analysis to be conducted in native cortical surface space.

The resulting outputs were then converted into a MATLAB-compatible format for further analysis using the *samsrf_mgh2srf* command from the SamSrf toolbox (version 7.13; https://github.com/samsrf). pRF estimates were computed by fitting a 2D Gaussian model to the data. As a result, three pRF parameters for each vertex on the cortical surface were computed: the x- and y-coordinates of the pRF centre and the pRF size (σ). Eccentricity and polar angle maps were computed from the x- and y-coordinates.

To model population receptive fields, a symmetric bivariate Gaussian model was used, with mean (x, y) representing the preferred retinotopic location, and standard deviation (σ) representing pRF size. To identify the pRF model parameters (x, y, σ) that best predict the measured time series, a two-stage fitting procedure was employed. In a coarse fitting step, data were smoothed along the cortical surface (2D Gaussian FWHM kernel of 5 mm) and a grid-search approach was used to identify model parameters that maximise the Pearson correlation between observed data and the pRF model’s predicted time course. Vertices with R^2^ > 0.05 were entered as starting values in a fine-fitting step, which used MATLAB’s *fminsearch* function to identify parameters that minimised the squared residual deviations between the model and unsmoothed data. Finally, X and Y position estimates were converted to eccentricity (distance from fixation) and polar angle. After that, each map was visually inspected for quality control before group-averaging (see *Technical Validation*).

After estimating pRF parameters on each participant’s native cortical surface, the resulting maps were projected to a common reference space to enable group-level analysis. This was achieved by resampling each individual’s maps from their native cortical surface onto the *fsaverage* template surface provided by FreeSurfer (*Native2TemplateMap* function in SamSrf v7.13). The resampled maps were then averaged vertex-wise across participants on the *fsaverage* mesh, resulting in group-level retinotopic maps for each parameter: eccentricity, polar angle, and pRF size.

## Data Records

Information and anatomical data that could be used to identify participants have been removed from all records. The resulting dataset is available on the OpenNeuro.org platform (https://openneuro.org/datasets/ds006642). The scripts used for data processing are available on the GitHub repository (https://github.com/levchenkoegor/movieproject2).

### Questionnaires

**Location** sourcedata/Questionnaire_participants_labels.csv

**File format** comma-separated value (CSV)

Participants’ responses to a battery of questionnaires (see *Questionnaires* under *Tasks* section) in a comma-separated value (CSV) file. Data is structured as one line per participant with all questions and test items as columns.

### Cognitive tasks

**Location** sourcedata/Cognitron_assessment_participants_cleaned.tsv

**File format** Tab-separated values (TSV)

Participants’ responses to a battery of cognitive tasks (see *Cognitive Tasks* under *Tasks* section) in a TSV file. The file contains the *sub-id* identifier column and *data* with all responses organised in a JSON format. The table is structured as one line per task (16 rows per participant).

### Anatomical MRI

**Location** sub-<ID>/ses-<SES_ID>/anat/sub-<ID>_ses-<SES_ID>_T1w.nii.gz

**Session** 001, 002

**File format** NIfTI, gzip-compressed

**Sequence protocol**

sub-<ID>/ses-00<SES_ID>/anat/sub-<ID>_ses-<SES_ID>_T1w.json

The anatomical scan was acquired at the end of the first session (5-min MPRAGE) and at the beginning of *Session 2* (2-min MPRAGE). Both raw anatomical images were defaced. They are available as a 3D image file, stored as sub-<ID>_ses-<SES_ID>_T1w.nii.gz. The sequence protocol file accompanies anatomical images in the same folder, stored in JavaScript Object Notation (JSON) file format.

### Functional MRI

**Location**

sub-<ID>/ses-<SES_ID>/func/sub-<ID>_ses-<SES_ID>_task-<TASK_NAME>_run-< RUN_ID>_bold.nii.gz

**Session** 001, 002

**Task-name** backtothefuture, somatotopy, retinotopy, tonotopy

**Run** 001, 002, 003

**File format** NIfTI, gzip-compressed

**Sequence protocol**

sub-<ID>/ses-<SES_ID>/func/sub-<ID>_ses-<SES_ID>_task-<TASK_NAME>_run-<RUN_ID>_bold.json

Raw fMRI data is available as individual time series files for each task run.

**Location**

sub-<ID>/ses-<SES_ID>/fmap/sub-<ID>_ses-<SES_ID>_acq-<TASK_NAME>_dir-PA_run-<RUN_ID>_epi.nii.gz

**Session** 001, 002

**Task-name** func, somatotopy, retinotopy, tonotopy

**Run** 001, 002, 003

**File format** NIfTI, gzip-compressed

**Sequence protocol**

sub-<ID>/ses-<SES_ID>/fmap/sub-<ID>_ses-<SES_ID>_acq-<TASK_NAME>_dir-PA_run-<RUN_ID>_epi.nii.json

The phase reverse encoding (P >> A) scan was acquired for each task and is available as an individual time series. The *func* task-name refers to the movie-watching task (*backtothefuture* in all other files). The sequence protocol file accompanies functional images in the same folder, stored in a JSON file.

### Physiological recordings

**Location**

sourcedata/sub-<ID>/ses-<SES_ID>/func/sub-<ID>_ses-<SES_ID>_run-<RUN_ID>_task-<TASK_NAME>_physioPULS.tsv

**Session** 001, 002

**Task-name** backtothefuture, retinotopy, tonotopy

**Run** 001, 002, 003

**File format** TSV

**Information file**

sub-<ID>_ses-<SES_ID>_run-<RUN_ID>_task-<TASK_NAME>_physioInfo.tsv

Raw pulse data (acquired with a pulse oximetry sensor) is available as individual time series files in TSV format.

### Eye tracker recordings

**Location**

sourcedata/sub-<ID>/ses-<SES_ID>/func/sub-<SUB_ID>_ses-<SES_ID>_run-<RUN_ID>_task-<TASK_NAME>_eyelinkraw.<FORMAT>.gz

**Session** 001, 002

**Task-name** backtothefuture, retinotopy

**Run** 001, 002, 003

**File format** EDF or ASC, gzip-compressed

Raw eye-tracker data is available in two formats: EDF and ASC. Raw eye-tracker EDF files were converted to ASCII files using the *edf2asc* tool (EDF2ASC version 4.2.1197.0) from the Eyelink Developers Kit. The ASCII files contain messages (‘MSG’) flags to align the EPI and gaze positions data. ‘MOVIE_START’ indicates the beginning of the movie, ‘FRAMENUMBER_*” indicates the number of frames shown on the screen and “PULSE_*” indicates the number of pulses received from the scanner. The snippet of the ASCII file with messages is provided below:

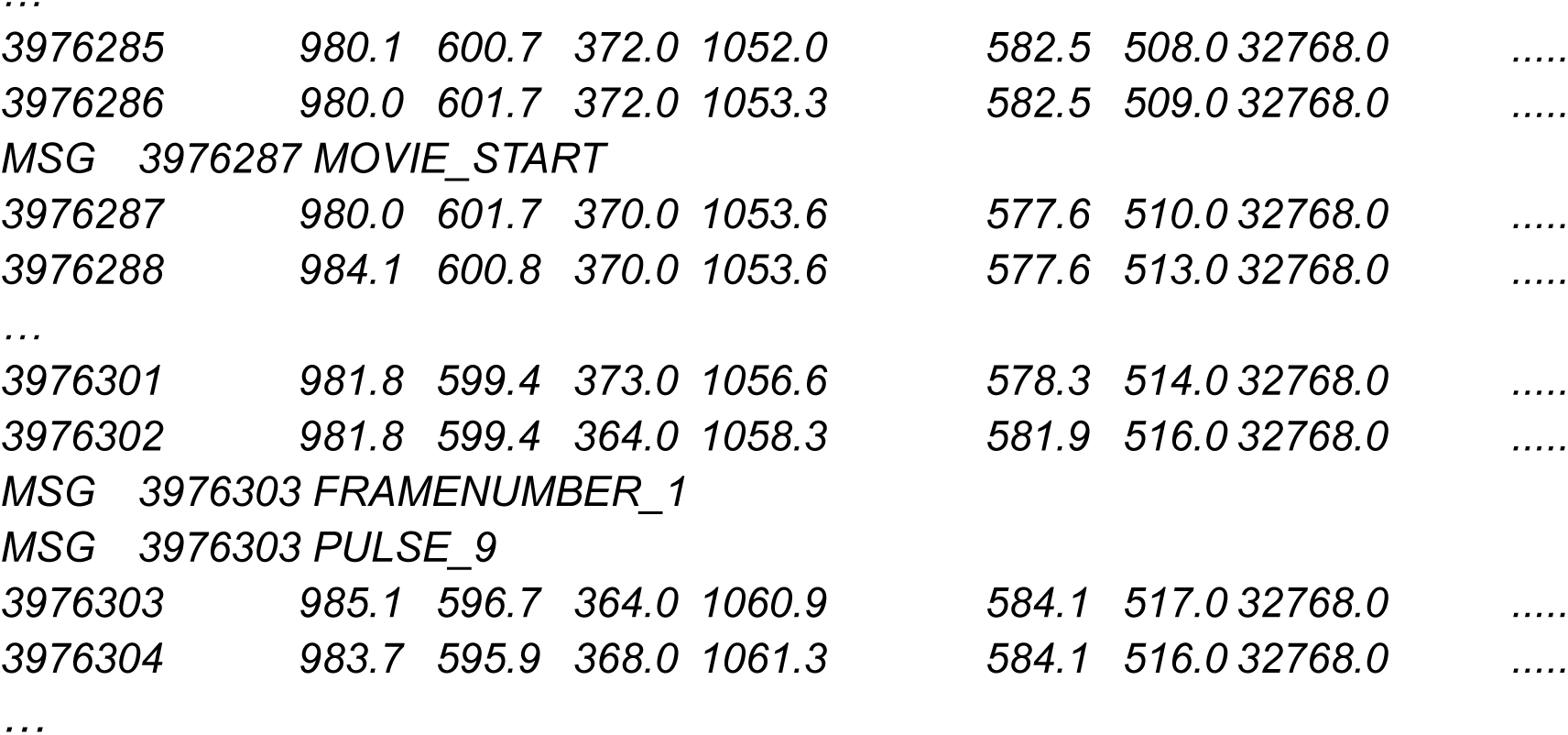

In the current snippet, there are events in the ‘MSG’ lines. In a similar way, the retinotopic mapping eye tracker ASCII files have ‘MSG’ for each volume. ‘VOLUME_*” indicates the number of pulses received from the scanner.

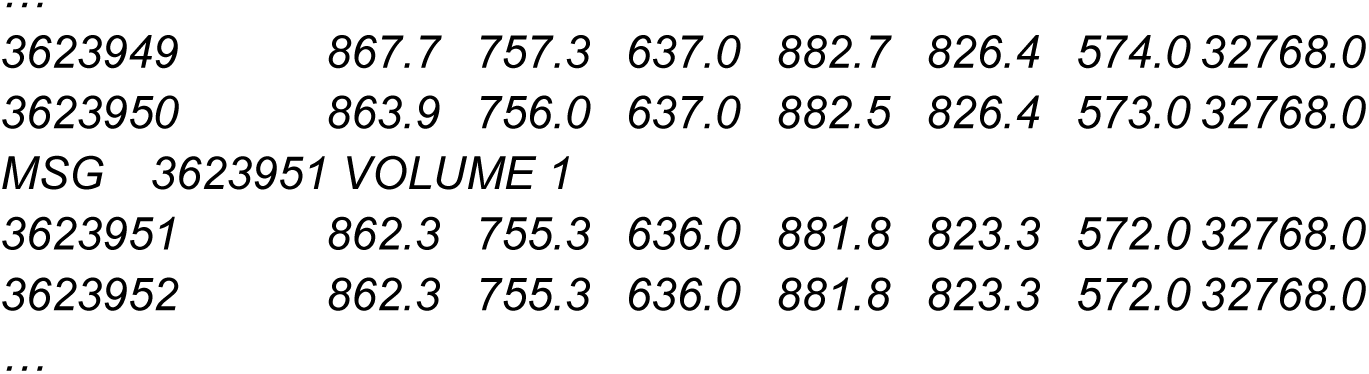

In both cases, the messages (‘MSG’) received from the MATLAB script make it possible to align the eye-tracker and EPI data precisely.

### Freesurfer outputs

**Location** derivatives/freesurfer/sub-<ID>

The standard Freesurfer outputs are obtained after running the *recon-all* command for each participant separately. In each folder, there is a SUMA folder with all the outputs from the *SUMA_Make_Spec_FS* command. The pRF maps are stored under the *retinotopy* folder.

### SSwarper outputs

**Location** derivatives/sub-<ID>/SSwarper

The standard SSwarper outputs after running the *SSwarper* AFNI command for each participant separately. This procedure both skull-strips the anatomical volume and computes the nonlinear warp to standard space.

### Preprocessed functional MRI

**Location**

derivatives/sub-<ID>/<TASK_NAME>/sub-<ID>_task-<TASK_NAME>_run-<RUN_ID>_preproc.nii.gz

**Task-name** backtothefuture, somatotopy, tonotopy, retinotopy

**Run** 001, 002, 003

The preprocessed fMRI data after running the AFNI processing script (*afni_proc.py*). The data is available as individual time series files for each task run. The preprocessing script for each task can be found in the GitHub repository.

## Technical Validation

### Evaluation of head motion control

The head motion was evaluated using the framewise displacement (FD) metric, which measures instantaneous head motion by comparing the motion between the current and previous volumes. The FD values were calculated using standard AFNI functionality (*3dvolreg* function). *Figure 4* shows the overall distribution of FD values for each task and run or condition. The distributions (first column) are highly skewed toward lower FD values, with the majority concentrated below 0.2 mm. A dashed red line marks the 95th percentile of the FD values at 0.2, 0.24, 0.2 and 0.25 mm for *backtothefuture*, *somatotopy*, *retinotopy* and *tonotopy* tasks respectively, indicating that 95% of the data exhibits minimal motion. Violin plots (second column) evaluate FD on a run or condition-specific level. Across all runs and conditions, FD distributions remain consistent, with median values below 0.2 mm and most of the values below 0.3 mm (for all tasks). These results underscore that head motion was well-controlled throughout the study, minimising potential motion-related artefacts in the data.

**Figure 2.**
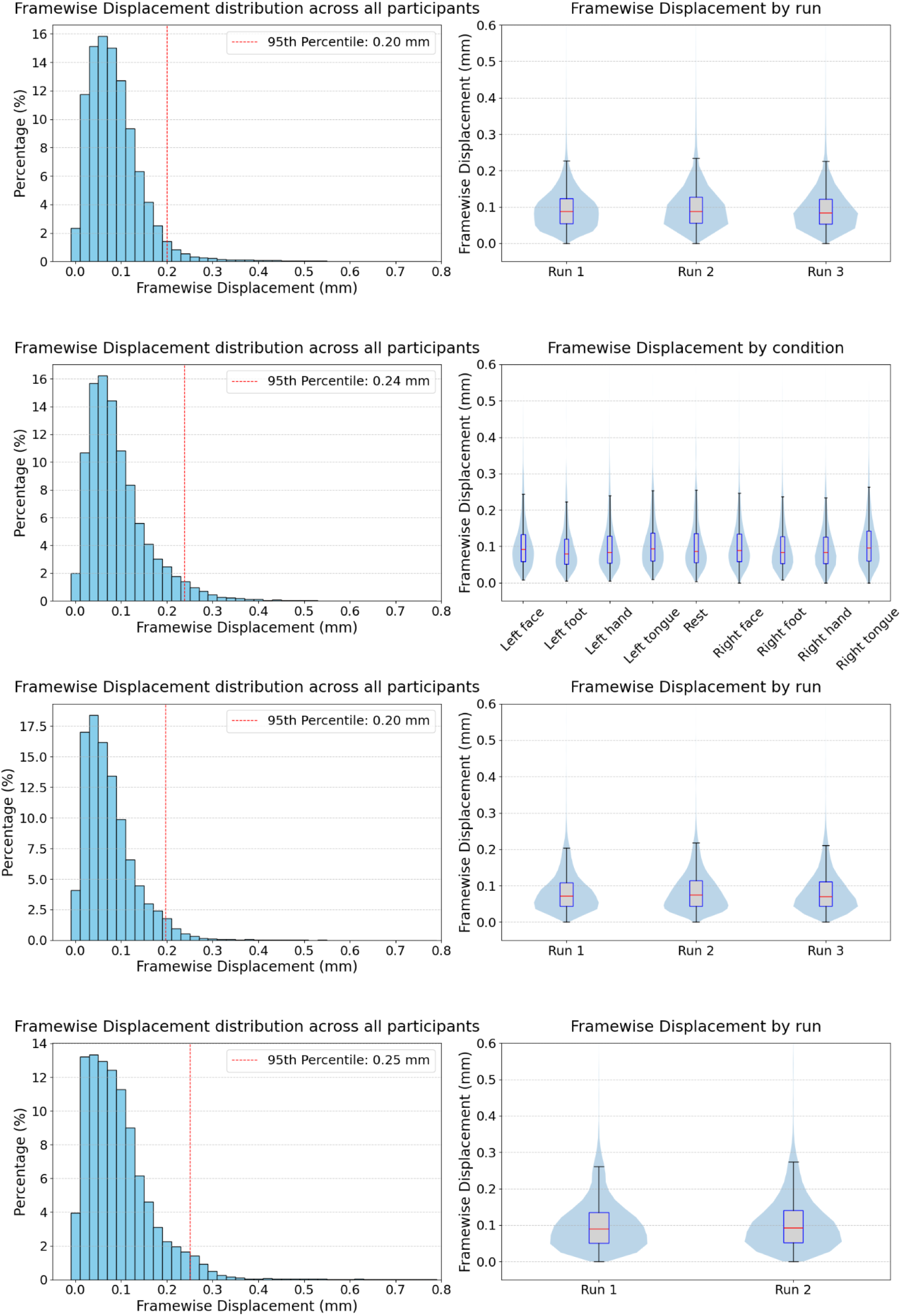
Framewise displacement distributions across all participants for each task and run. The figure shows the distributions for each task separately by row: backtothefuture, somatotopy, retinotopy, and tonotopy. The first column depicts the distribution of FD across all participants and TRs. The second column displays violin plots, and overlaid boxplots show the distribution of FD for each run or condition across all participants. Each violin plot represents the density of FD values, with the boxplot inside showing the interquartile range (IQR), the median, and whiskers (representing the minimum and maximum values within 1.5×IQR).

Table 3 shows descriptive statistics for head motion parameters across each task and run. It reports the mean, median, and maximum values for three rotational (Roll, Pitch, Yaw; in degrees) and three translational components (Superior-Inferior (dS), Left-Right (dL), Posterior-Anterior (dP); in millimetres). Overall, the mean displacements for both rotational and translational motion remained within ±0.1 degrees and millimetres, respectively. Mean maximum values for all motion parameters across all tasks were below 0.5, except the *backtothefuture* task, where maximum displacements in pitch and dS approached 1.0 degree/millimetre across all runs. Despite minor variations between tasks and runs, overall motion levels were low, suggesting minimal motion-related artefacts in subsequent analyses.

**Table 3.** Descriptive statistics for head motion parameters across each task and run. Each cell represents the mean, median, or maximum value of framewise displacement per run. Tasks include backtothefuture, somatotopy, retinotopy, and tonotopy, with up to three runs per task.

|  |  | Mean |  |  |  |  |  | Mean maximum |  |  |  |  |  |
| --- | --- | --- | --- | --- | --- | --- | --- | --- | --- | --- | --- | --- | --- |
| Task | Run | Roll | Pitch | Yaw | dS | dL | dP | Roll | Pitch | Yaw | dS | dL | dP |
| backtothefuture | 1 | -0.01 | -0.01 | 0.02 | -0.02 | -0.08 | -0.03 | 0.25 | 0.63 | 0.36 | 0.6 | 0.19 | 0.45 |
|  | 2 | -0.05 | 0.09 | 0.04 | -0.05 | -0.03 | -0.03 | 0.28 | 0.92 | 0.44 | 0.7 | 0.25 | 0.48 |
|  | 3 | -0.02 | 0 | -0.03 | 0.02 | -0.06 | -0.04 | 0.3 | 0.84 | 0.45 | 0.94 | 0.28 | 0.44 |
| somatotopy | 1 | -0.02 | 0.01 | -0.01 | 0.02 | -0.01 | -0.04 | 0.17 | 0.46 | 0.22 | 0.32 | 0.17 | 0.36 |
|  | 2 | -0.01 | 0 | 0.02 | 0 | -0.02 | -0.02 | 0.18 | 0.45 | 0.22 | 0.38 | 0.16 | 0.4 |
| retinotopy | 1 | -0.01 | 0.1 | -0.01 | -0.03 | -0.01 | -0.08 | 0.14 | 0.48 | 0.17 | 0.21 | 0.12 | 0.28 |
|  | 2 | 0 | 0.09 | 0.02 | 0.03 | 0.01 | -0.08 | 0.11 | 0.43 | 0.17 | 0.34 | 0.12 | 0.23 |
|  | 3 | -0.02 | 0.08 | 0.02 | 0.04 | -0.02 | -0.07 | 0.09 | 0.48 | 0.18 | 0.34 | 0.1 | 0.23 |
| tonotopy | 1 | -0.02 | 0.09 | -0.01 | 0 | 0 | -0.02 | 0.12 | 0.43 | 0.14 | 0.26 | 0.1 | 0.23 |
|  | 2 | 0.01 | 0 | 0.02 | 0.04 | 0.02 | -0.02 | 0.15 | 0.36 | 0.21 | 0.44 | 0.16 | 0.3 |

### Inter-subject correlation during movie-watching

The inter-subject correlation (ISC) analysis was conducted on all participants to demonstrate the timing alignment and the quality of functional MRI data during *backtothefuture* task. ISC is a data-driven method that quantifies shared temporal fluctuations in brain activity across participants (Nastase et al., 2019). It is commonly used to assess synchronisation of neural responses during naturalistic paradigms. Six participants were excluded from this analysis (see *Usage Notes* for details).

The ISC values were calculated using AFNI tools (*3dTcorrelate* and *3dISC*). Pairwise ISCs were computed for every unique pair of participants using Pearson correlation and Fisher z-transformation. Once all pairwise ISC maps were generated, a group-level statistical model was estimated using mixed-effects modelling that treated both participants in each pair as random intercepts. To assess statistical significance, t-statistics were subsequently corrected for multiple comparisons using the False Discovery Rate (FDR) procedure at a threshold of q < 0.001. The ISC values are shown in *Figure 3* below.

**Figure 3.**
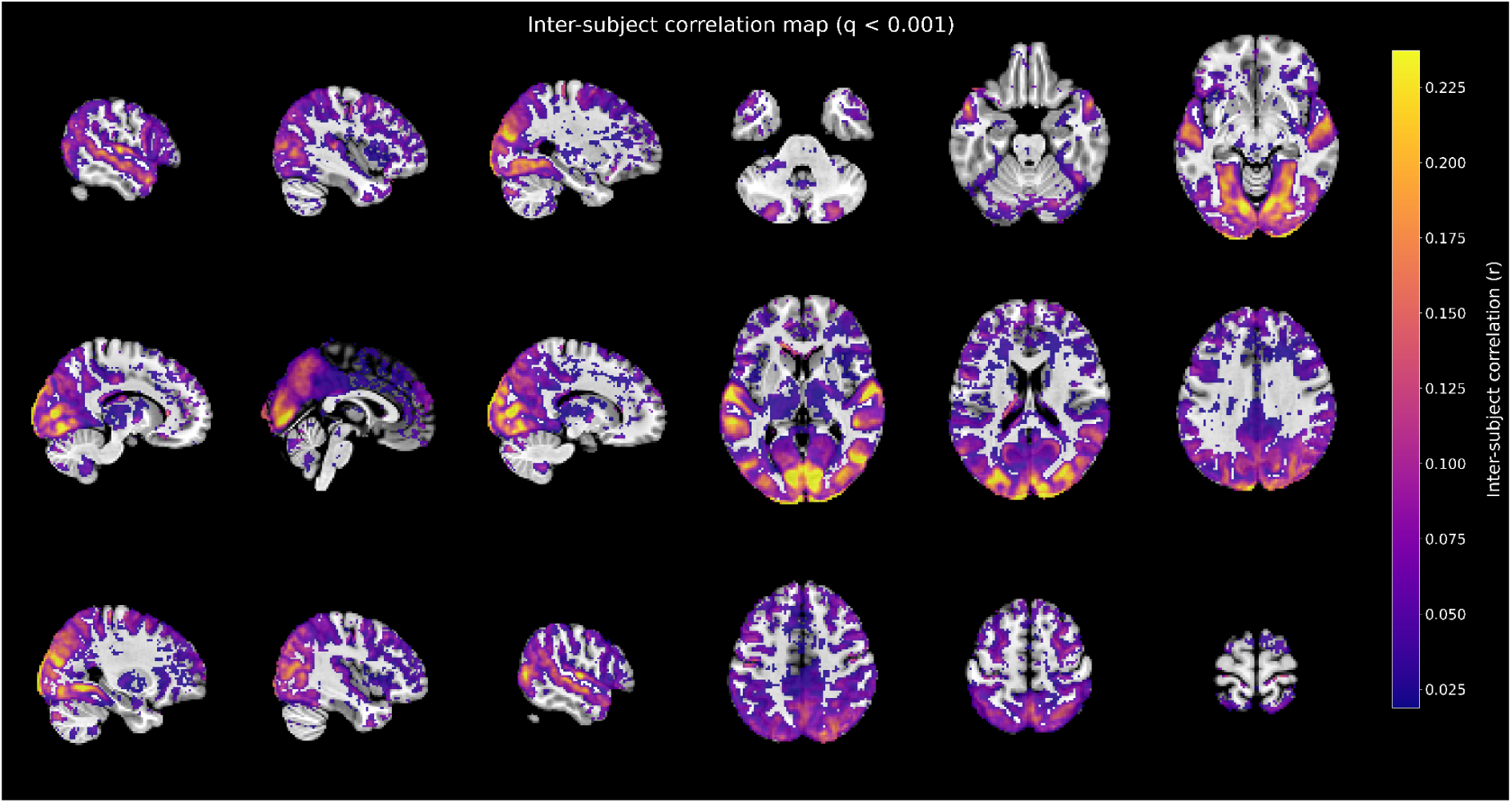
Inter-subject correlation map. Voxelwise inter-subject correlation (ISC) during Back to the Future viewing, computed across 34 participants after excluding six with alignment issues (see Usage Notes). Statistical significance was assessed with voxelwise t-tests and controlled for multiple comparisons using the False Discovery Rate (FDR) procedure at q < 0.001 (two-tailed). No additional cluster-size threshold was applied.

High ISC values across early sensory cortices further indicate accurate temporal alignment across participants. This suggests that both stimulus presentation and preprocessing preserved the temporal structure of the data, ensuring that stimulus-locked neural responses were synchronised across individuals. The resulting ISC maps show robust inter-subject synchronisation across widespread cortical regions, including primary auditory and visual cortices, as well as posterior medial areas (precuneus, posterior cingulate). These regions are consistent with previous findings in the literature and reflect reliable, stimulus-locked processing during naturalistic movie-watching tasks (Aliko et al., 2020; Hasson et al., 2004; Lerner et al., 2011).

### Eye-tracker quality control

Each participant underwent a standard 9-point calibration and validation procedure prior to each run of the *backtothefuture* task. To quantify calibration quality, the average and maximum errors for each eye and each run for all participants were extracted from the corresponding Eyelink ASCII files. The table with average and maximum errors per participant, run, and eye is available in the *Supplementary Materials*.

Following SR Research guidelines, calibration accuracy was classified as *GOOD* when the average error was below 1.0° and the maximum error was below 1.5°, *FAIR* when the average error ranged between 1.0° and 1.5° or the maximum error between 1.5° and 2.0°, and *POOR* when the average error exceeded 1.5° or the maximum error exceeded 2.0° (https://www.sr-research.com/support/showthread.php?tid=244). Across all calibration entries (N = 234), the mean average error was 0.46° (SD = 0.19°), and the mean maximum error was 1.03° (SD = 0.82°). Based on the classification criteria from the guidelines, 212 calibrations (90.6%) were labelled as *GOOD*, 13 (5.6%) as *FAIR*, and 9 (3.8%) as *POOR*.

Since we used a binocular setup, we adopted the following approach for run-level quality classification: if at least one eye showed *GOOD* calibration, the run was considered *GOOD*. Under this criterion, only 3 runs were labelled *POOR* and 2 as *FAIR*, while all remaining runs met the *GOOD* threshold for at least one eye (and in most cases, for both eyes). These results indicate a high level of calibration accuracy and confirm the overall quality of the eye-tracking data.

### Somatotopic mapping

To assess the quality and specificity of the somatotopic mapping, we first estimated participant-level beta coefficients using a general linear model (implemented in AFNI’s *3dDeconvolve* function with *dmUBLOCK(1)* parameter), modelling each body movement separately. Individual results were visually inspected to ensure expected activation patterns and data quality. We examined group-level activation maps for three contrasts between different body part movements: Face vs Feet, Face vs Hand, and Hand vs Tongue. For each participant, condition contrasts (Face vs Feet, Face vs Hand, and Hand vs Tongue) were computed within *3dDeconvolve*, resulting in subject-level contrast beta maps. At the group level, these contrast betas were then entered into a one-sample t-test to assess consistency across participants. As shown in *Figure 4*, the three contrasts show the activation maps aligned with the canonical somatotopic organisation along the precentral and postcentral gyri. Specifically, face-related activity was localised to the inferior-lateral portion of the sensorimotor cortex. Tongue-related activations were found ventrally, with strong effects in lateral sensorimotor regions and inferior pre-/postcentral areas. In contrast, feet-related activity was located dorsomedially, while hand activations occupied intermediate dorsolateral regions of the central sulcus. The results are consistent with the known motor and somatosensory homunculus, indicating both the spatial specificity and sensitivity of the task and data acquisition.

**Figure 4.**
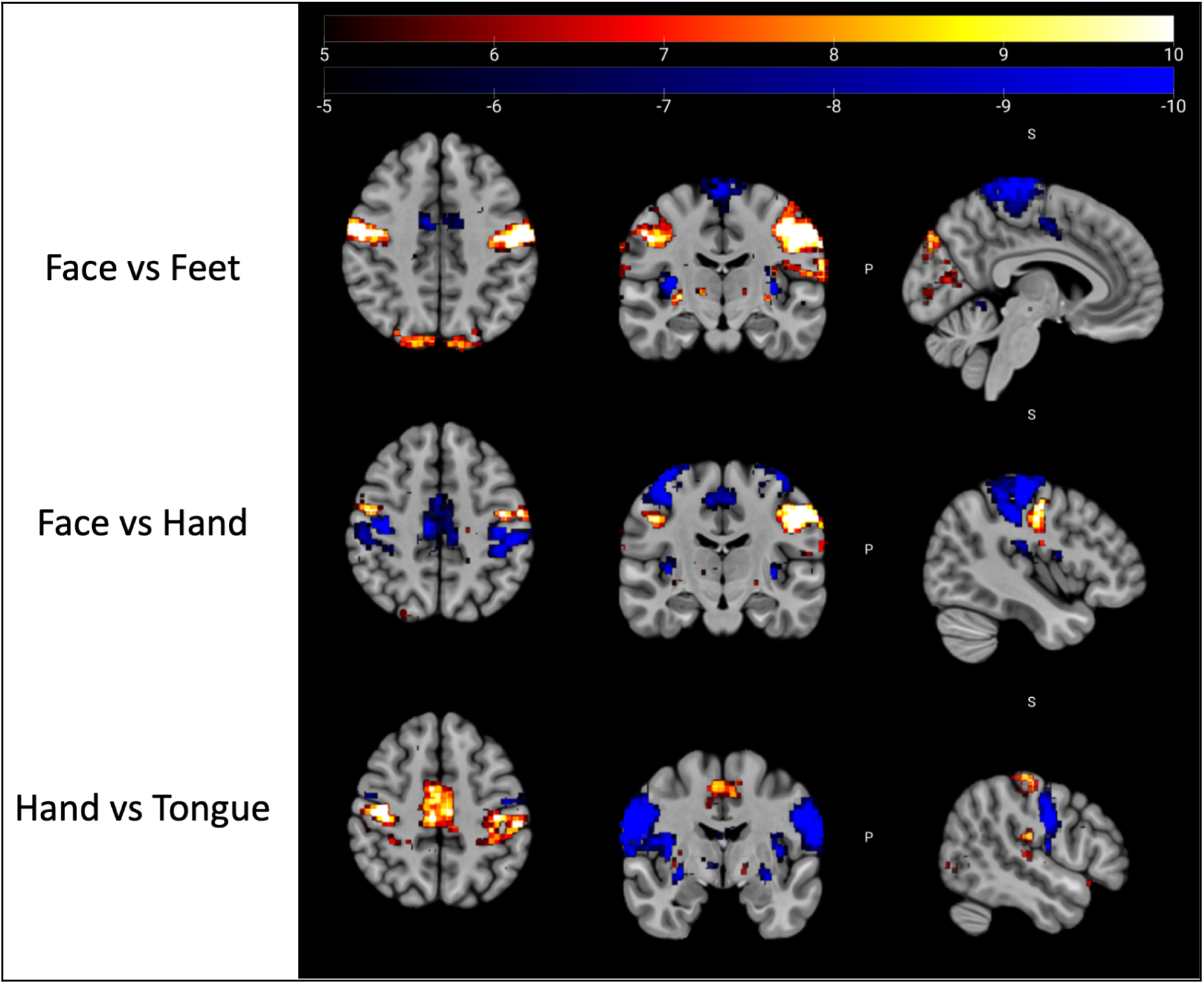
Somatotopic mapping. Group-level statistical activation maps are displayed for three pairwise contrasts: Face vs Feet, Face vs Hand, and Hand vs Tongue, each rendered on a standard MNI space in axial, coronal and sagittal planes. Colours represent voxel-wise t-values, thresholded at t > 5, corresponding to p < 0.00001. Red to white colour indicates regions showing significantly greater activation for the first body part in each contrast, while blue to dark blue colour indicates greater activation for the second body part.

### Retinotopic mapping

To evaluate the quality of the retinotopic mapping, we first estimated pRF parameters for each participant separately. Each participant’s eccentricity and polar angle maps were visually inspected for anatomical consistency, smoothness of topographic gradients, and the presence of expected retinotopic features. This evaluation was qualitative, focusing on whether the maps displayed coherent and interpretable organisation across early visual areas.

Participants 07, 17, and 29 were excluded from the group-level analysis due to poor retinotopic map quality. Although data were collected for these individuals, their maps lacked coherent topographic organisation, showing either excessive noise or fragmented patterns without the expected polar angle reversals or eccentricity gradients. As a result, these maps could not be reliably interpreted and were omitted from the mean maps across all participants.

To show the quality of the *retinotopy* task, the maps were averaged across all the remaining participants to get a grand average map (*Fig. 5*). Polar angle shows the well-known pattern along the calcarine sulcus and around the occipital cortex. In both hemispheres, polar angle values smoothly transition from the horizontal meridians (coded in green) to upper and lower vertical representations (coded in blue and red, respectively), consistent with known topography in areas V1, V2, and V3. Eccentricity maps show a clear central-to-peripheral gradient, with foveal representations located at the occipital pole (coded in blue) and peripheral regions exhibited more anteriorly (green). The resulting maps show well-organised and expected representations of both polar angle and eccentricity on inflated cortical surfaces of each hemisphere.

**Figure 5.**
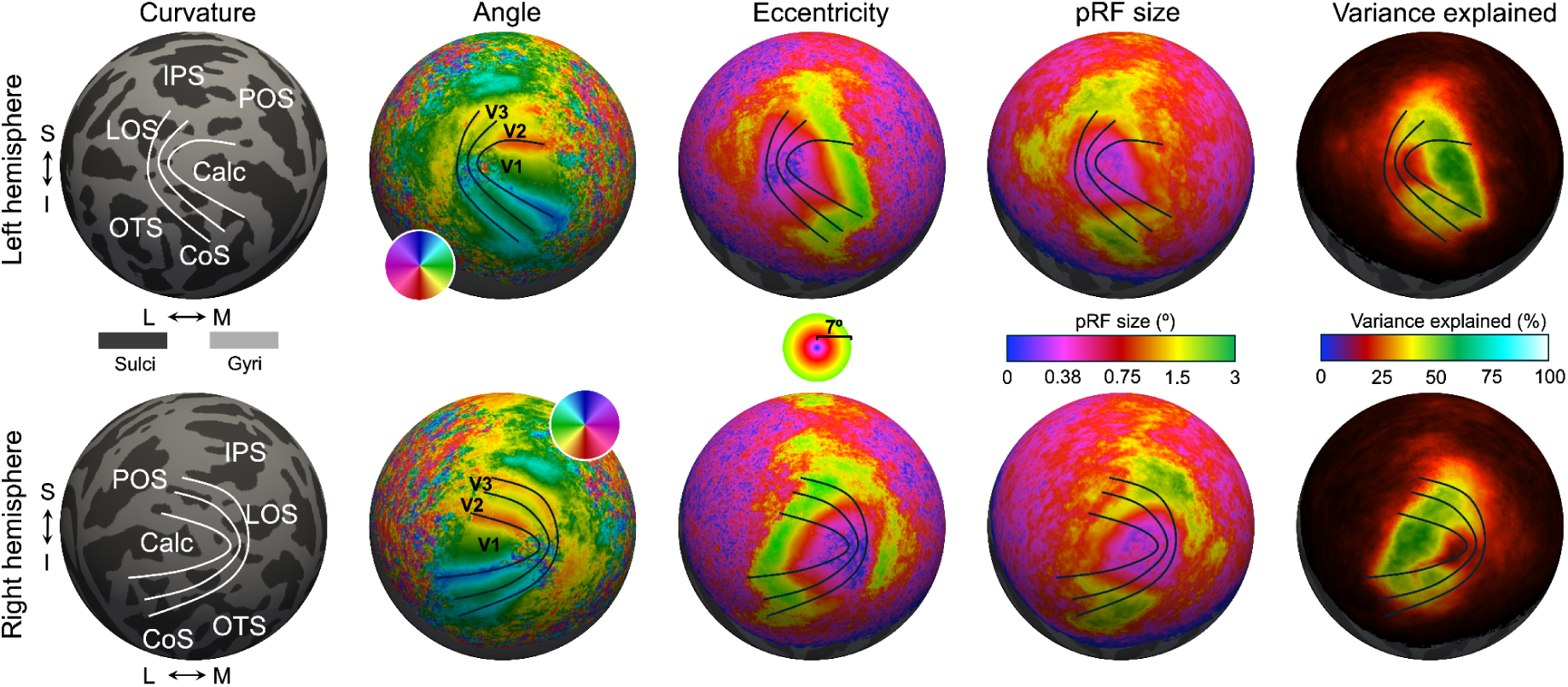
Retinotopic mapping. pRF results for the left and right hemispheres are displayed on inflated cortical surface reconstructions. Columns from left to right show: cortical curvature (dark gray, sulci; light gray, gyri) with anatomical landmarks labeled (IPS, intraparietal sulcus; POS, parieto-occipital sulcus; LOS, lateral occipital sulcus; Calc, calcarine sulcus; OTS, occipito-temporal sulcus; CoS, collateral sulcus); polar angle maps indicating visual field representation (color wheel inset); eccentricity maps showing distance from fixation (scale inset, center 0°, periphery 7°); estimated pRF size (in degrees of visual angle, color scale inset); and variance explained by the pRF model (in %, color scale inset). Black lines delineate visual area boundaries (V1–V3). Rows correspond to the left hemisphere (top) and right hemisphere (bottom).

## Usage notes

The NNDb-3T+ dataset provides a uniquely rich multimodal resource for studying the brain under naturalistic conditions and individual differences across multiple domains (somatotopy, retinotopy and tonotopy). The dataset combines high-quality fMRI data from a full-length movie-watching paradigm and somatotopy, retinotopy, and tonotopy tasks, eye-tracking data, physiological recordings, and extensive behavioural measures.

### Practical considerations

Several participants were excluded from inter-subject correlation analyses due to interruptions or inconsistencies during the movie task. These participants should be carefully considered for any time alignment-sensitive analysis and potentially excluded or adjusted. For many analyses, these participants could still be used.

**sub-01**: the scanner stopped near the end of run 2 which means that the last 152 TRs of the run were not acquired. Run 3 was collected with no issues.

**sub-02**: incorrect lengths of runs. The movie files were cut incorrectly: run 2 is missing the final 12 seconds, while run 3 includes those 12 seconds again (overlap with run 2). Perfect alignment can’t be guaranteed.

**sub-03**: scan stopped 55 seconds (37 TRs) before the end of run 2. Run 3 was collected with no issues.

**sub-05:** the anatomical MRI is missing due to technical reasons.

**sub-10**: movie stopped ∼15 minutes (511 TRs) before the end of run 3 due to participant scheduling conflict.

**sub-24** and **sub-36**: paused during run 1 using the implemented pause/resume functionality (see below). Run 1 is not recommended for time-sensitive analysis; run 2 and 3 were collected with no issues.

The presentation MATLAB script for *backtothefuture* task included a pause functionality for situations in which participants squeezed the alert ball, indicating a need for a break during the run, or if the movie needed to be stopped for any reason. If the movie was paused (by pressing ‘P’ on the keyboard), it froze on the last frame displayed and waited for the ‘R’ key to be pressed to resume. When ‘R’ was pressed, the movie rewound by 12 seconds and paused for eight spare TRs from the scanner. The operator adjusted the protocol to ensure the correct number of TRs for the remainder of the movie and resumed scanning when the participant was ready. This functionality was used with *sub-24* and *sub-36*, both of whom paused during the first part of the movie. In the first case, the participant initially needed a bathroom break but decided to finish the first run and use the bathroom during the break between runs. In the second case, the participant appeared sleepy, prompting the operator to stop the scanning and allow a short break inside the scanner. After five minutes, the scanning resumed. In both cases, the pause resulted in two files for the first run, labelled with the prefixes *‘beforepause’* and *‘afterpause’* to distinguish them.

The database includes both raw and preprocessed MRI data, with preprocessing pipelines designed to follow best practices for functional neuroimaging using AFNI, Freesurfer, MATLAB and Python. These pipelines are tailored to the specific structure and goals of each task. However, we emphasise that no single preprocessing strategy is optimal for all hypotheses or analytical approaches. Depending on the research question, users may wish to modify aspects of the pipeline such as spatial smoothing, alignment methods, filtering, nuisance regression, or motion censoring thresholds. We encourage users to consult the accompanying GitHub repository (see *Code availability*), which provides all preprocessing scripts, quality control reports, and logs. Reviewing these materials can help ensure reproducibility, clarify processing decisions, and guide adaptations for custom workflows.

### Limitations

Due to copyright restrictions, the full-length movie stimulus used in the backtothefuture task cannot be included in the dataset. However, the copy of the movie can be purchased with their unique Amazon Standard Identification Number (ASIN: B000BVK82I) or International/European Article Number (EAN: 5050582401288). Researchers can perform or apply timecoded annotations using publicly available tools or resources. Users with a legally obtained copy of the film can align annotations using the frame-level metadata and scanner trigger messages provided in the dataset. The exact same movie stimulus as in NNDb v1.0 was used, and all extracted annotations from that version are equally applicable to the current dataset.

In recent years, the rise of machine learning and computer vision algorithms has dramatically improved the quality, scalability, and diversity of annotations available for naturalistic stimuli. Tools like Neuroscout (https://neuroscout.org), built on pipelines such as *Pliers*, offer frame-level annotations for many popular films, including *Back to the Future*, covering features such as dialogue, visual objects, faces, scenes, and even emotional tone. These automated annotations can be easily adapted for use with NNDb-3T+. At the same time, it is important to note that while such annotations are a valuable resource, they are not a prerequisite for meaningful analysis of the database. Many approaches in naturalistic neuroimaging, such as inter-subject functional connectivity (ISFC) (Ren et al., 2017), dynamic functional connectivity (Hutchison et al., 2013), or model-free analyses of temporal reliability (Nastase et al., 2020), do not require any annotations. Thus, the availability of annotations enhances the versatility of the dataset, but the scope of potential analyses extends well beyond annotation-based methods.

Although high-quality annotations can be generated or sourced for *Back to the Future*, the fact that the database includes only one movie presents an inherent limitation. Even with frame-level labels for visual, auditory, or linguistic features, the number of unique events, scenes, or semantic contexts remains finite. This may limit the statistical power and generalizability of certain modelling approaches. Similarly, attempts to study rare or specific event types (e.g. non-verbal social interactions without dialogue, ambiguous language use, animal sounds) may be constrained by insufficient samples.

## Code availability

All the code is available on the GitHub repository: https://github.com/levchenkoegor/movieproject2/.

## Acknowledgements

All MRI scanning hours were donated by the Birkbeck/UCL Centre for NeuroImaging (BUCNI) as part of its commitment to open science resources. Pilot scanning, participant funds, and a PhD tuition and stipend fellowship to EL work were supported by the London Interdisciplinary Doctoral Programme training grant (Biotechnology and Biological Sciences Research Council BB/T008709/1). We would like to thank Letitia Schneider, Raha Razin, Oliver Josephs, Joerg Magerkurth, Winnie Yeh and other staff at BUCNI for their support. We thank Agata Czarnecka and the whole Cognitron team for providing the platform for behaviour data collection. JIS is supported by the Wellcome Leap.

## Competing interests

The authors declare no competing interests.

## Supplementary materials

### Figures illustrating the trial structure of cognitive tasks

**Figure 1.**
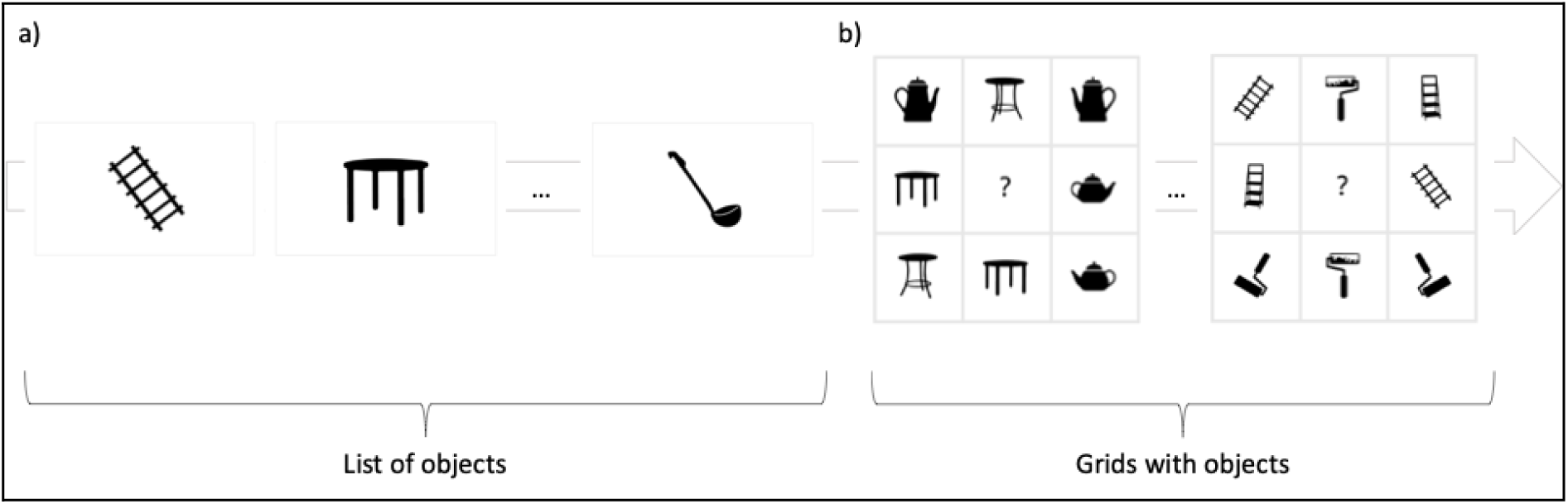
Schematic representation of the *Object Memory Immediate and Delayed* task. Panel *a)* shows a set of objects presented to the participant in the first part of the task. Panel *b)* shows the grids where participants needed to find and click on the object which was previously presented in the set.

**Figure 2.**
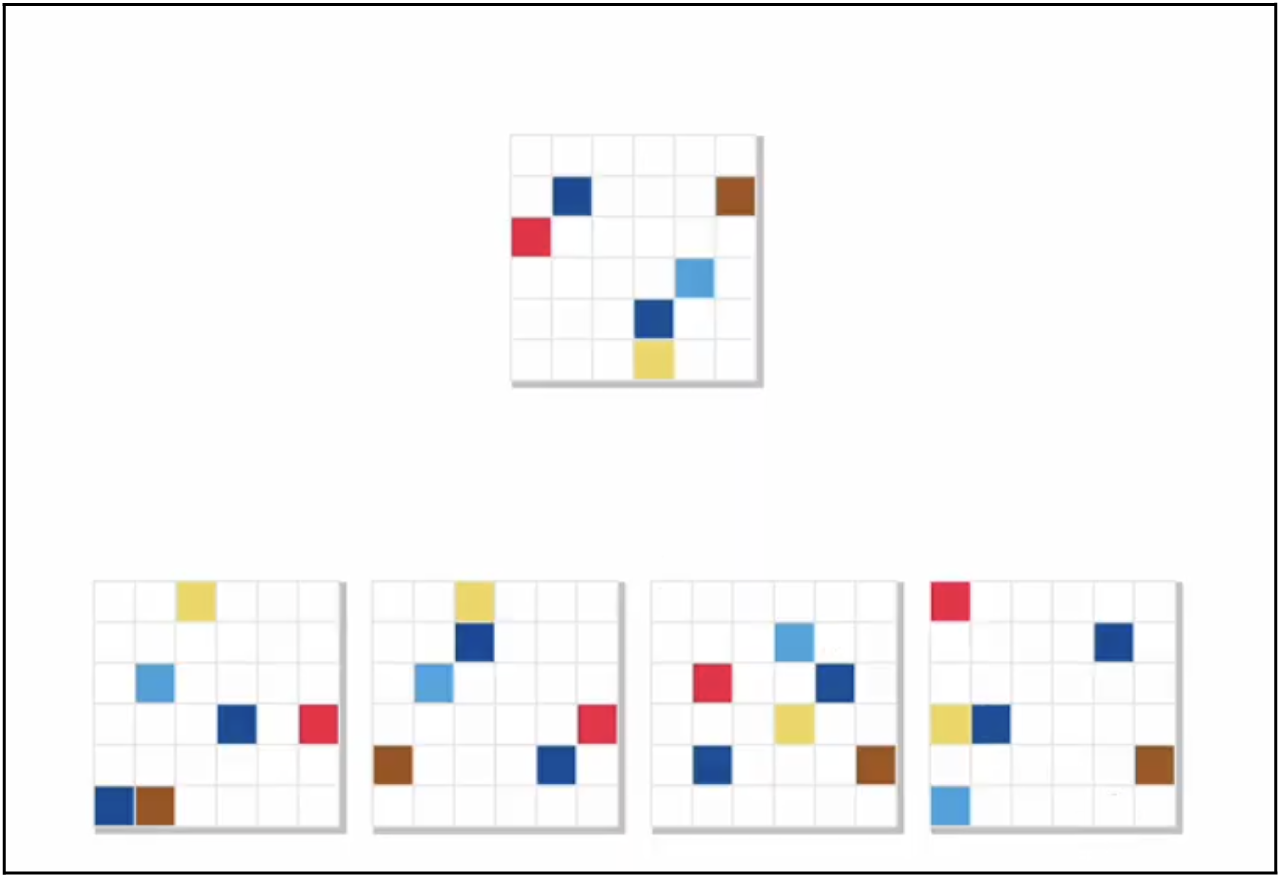
One trial for a *two-dimensional manipulation* task. The target grid is shown at the top, with four possible options displayed below. Participants were asked to identify which of the four grids represented a rotated transformation of the target grid. In the example shown, the correct answer is the second grid from the left, which corresponds to a 180-degree rotation of the target grid.

**Figure 3.**
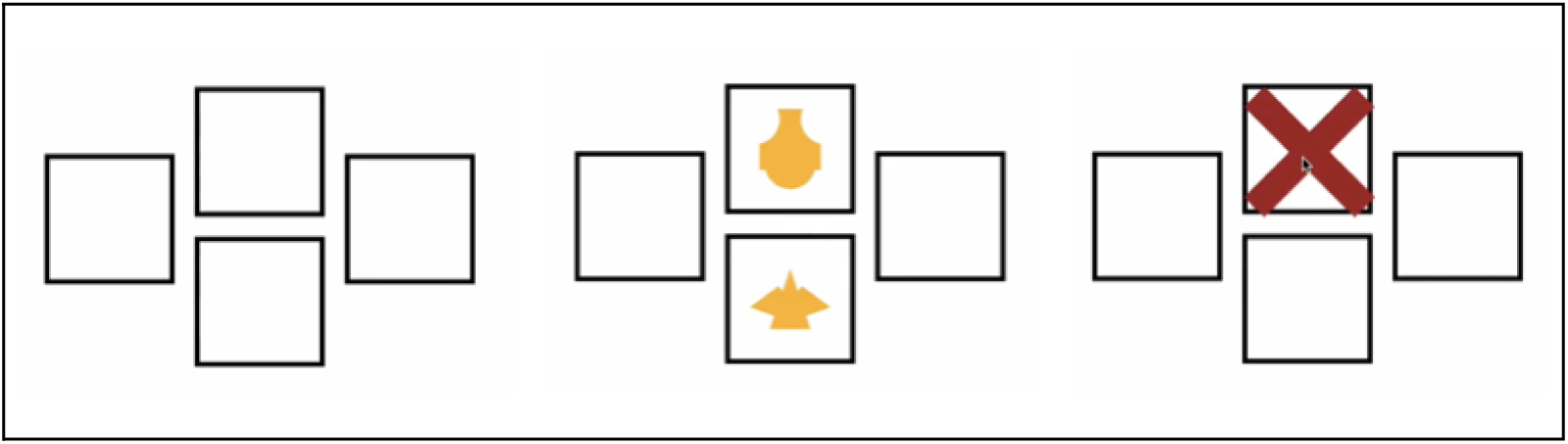
Example trial from the *Intra/Extra-Dimensional Set-Shifting (ID/ED)* task. Each trial consists of three consecutive screens: a blank screen briefly appears before stimulus onset; two compound stimuli are presented, requiring the participant to choose one. Feedback follows the participant’s response, indicating whether it was correct (e.g., a red cross for an incorrect response).

**Figure 4.**
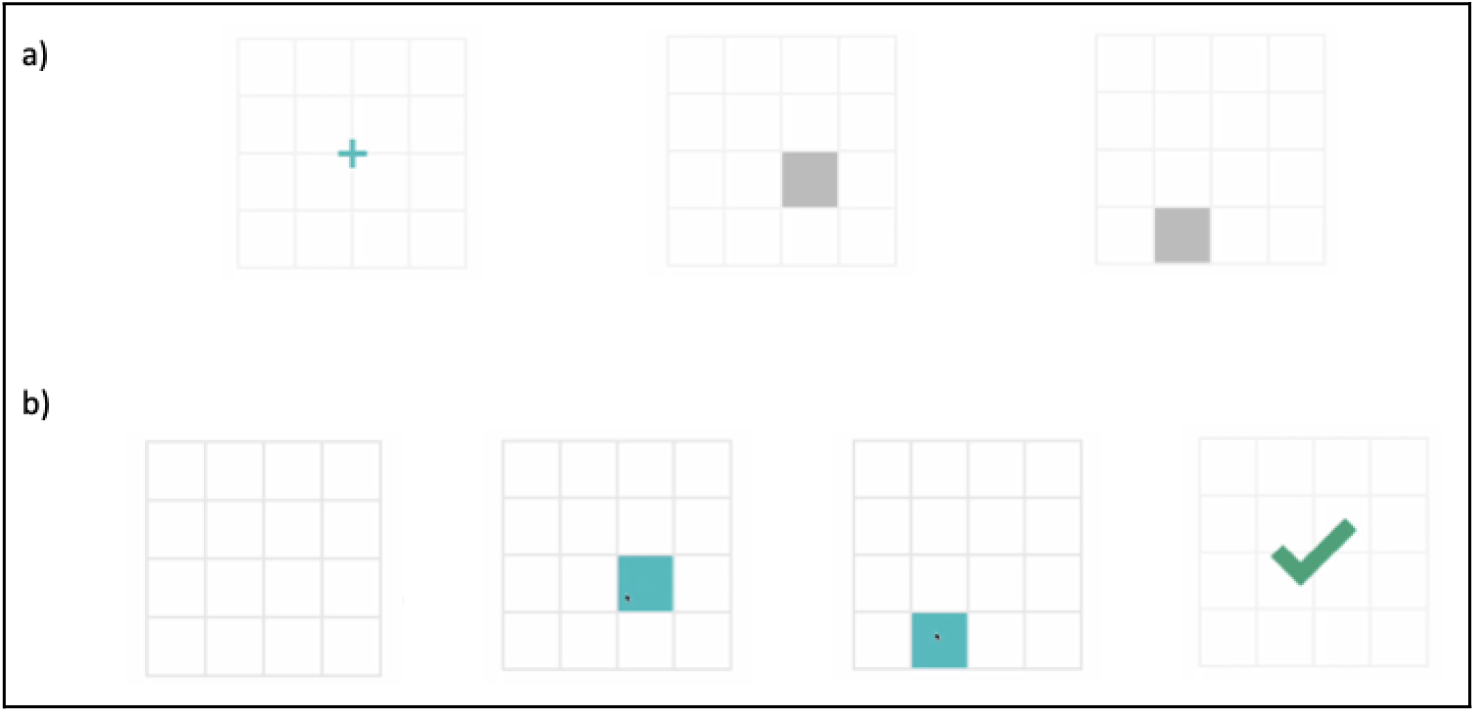
Trial of the *Spatial Span* task. Panel *a)* shows the presented sequence of squares lightening up (2 in the current example). Panel *b)* shows the empty grid and the correct response in this trial. The feedback was received after each trial.

**Figure 5.**
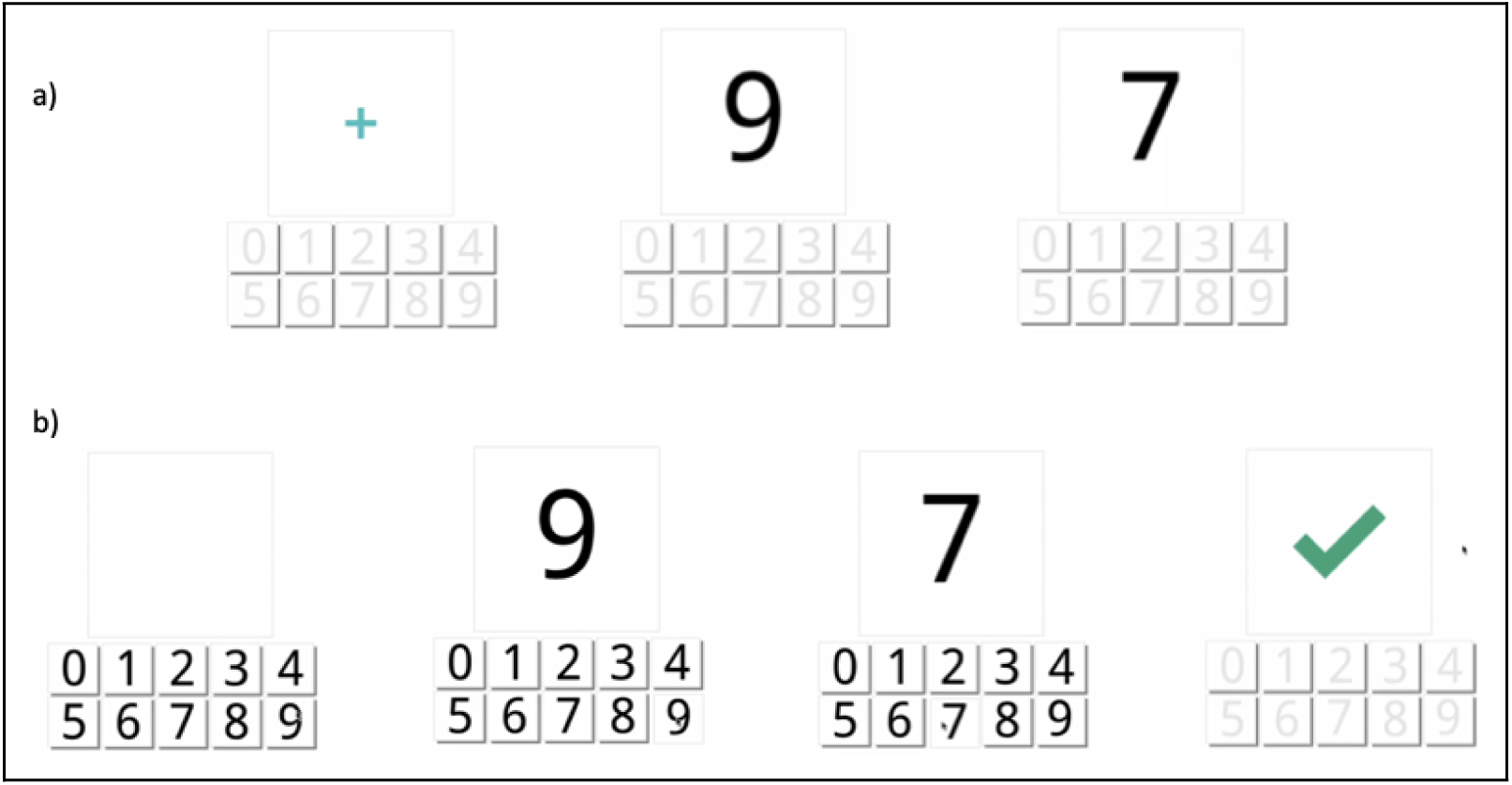
Trial of the *Digital Span* task. Panel *a)* shows a sequence of digits (2 in the current example). Panel *b)* shows the digital keyboard and the correct response in this trial. Feedback was received after each trial.

**Figure 6.**
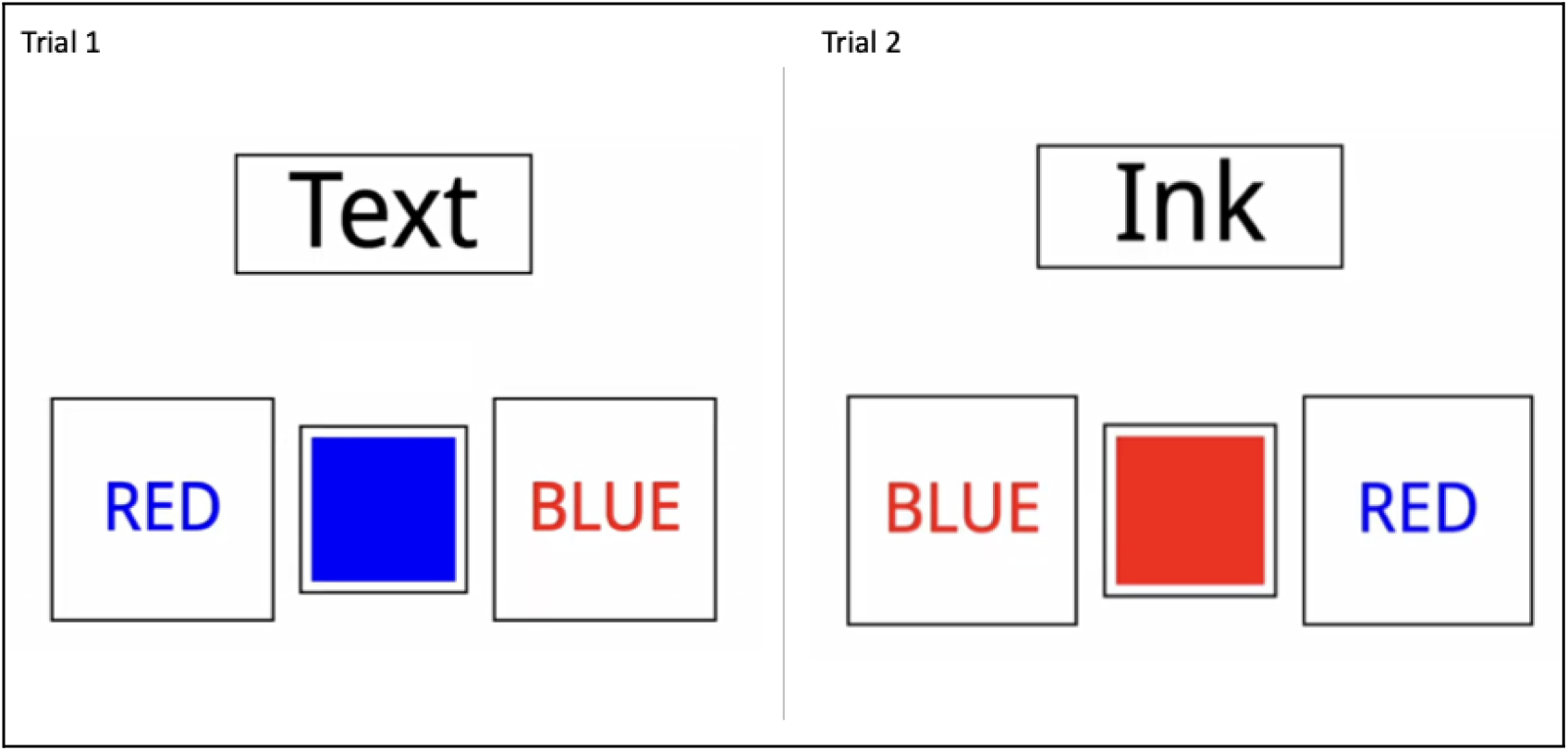
Two trials with two different conditions (Text or Ink) of the *Switching Stroop* task. Trial one (on the left side of the figure) instructed participants to click on the ‘RED’ box (written in blue colour). In the second trial (on the right side of the figure), the correct response was the ‘BLUE’ box (written in red colour).

**Figure 7.**
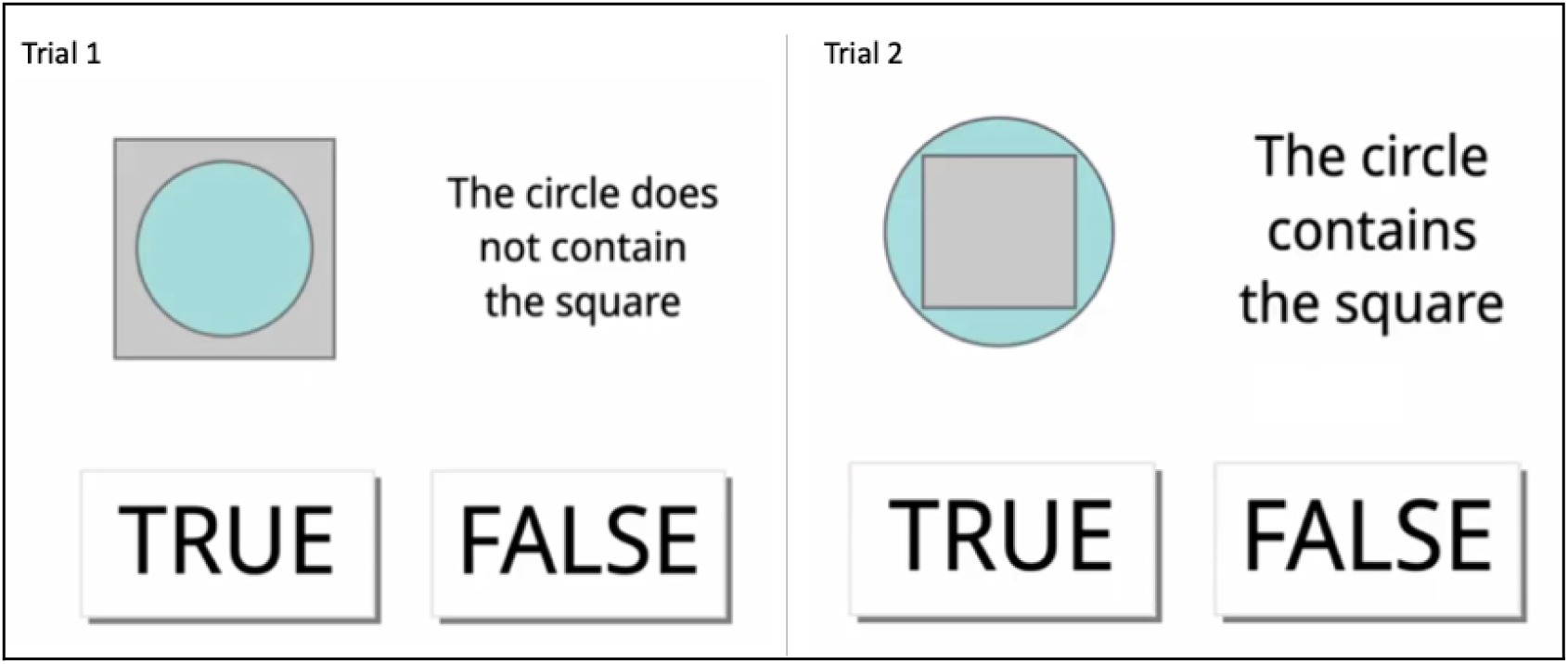
Two trials of the *Verbal Reasoning* task. The correct response in both trials is True.

**Figure 8.**
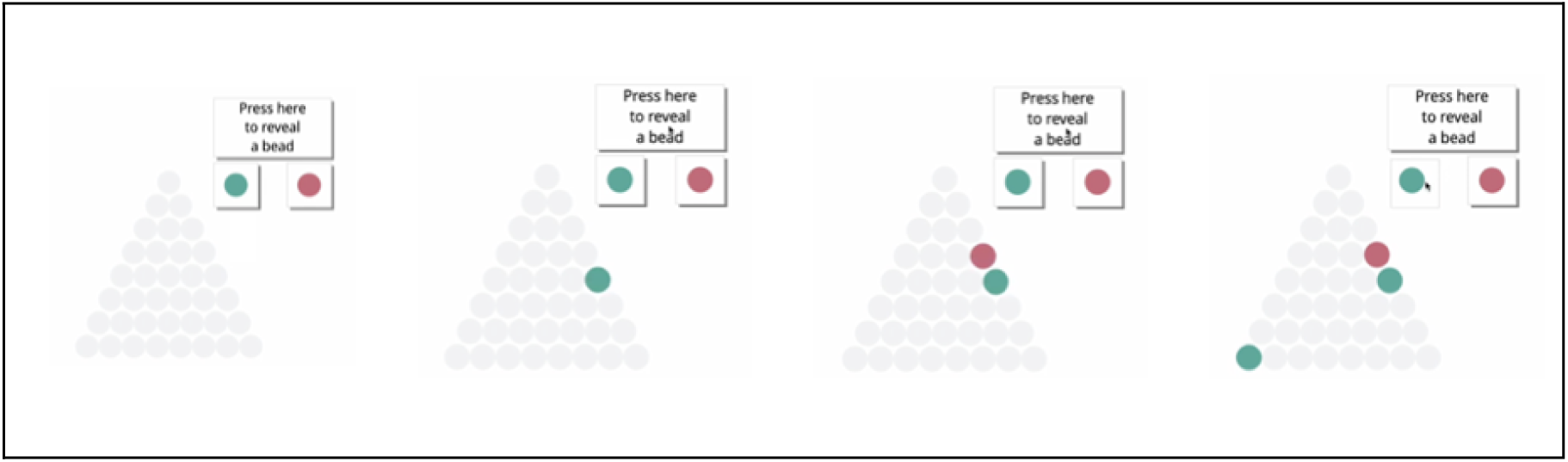
A trial for the *Beads* task. In this example, the participant revealed three beads and decided to pick green as the dominant colour in the jar. Feedback was shown after each trial.

**Figure 9.**
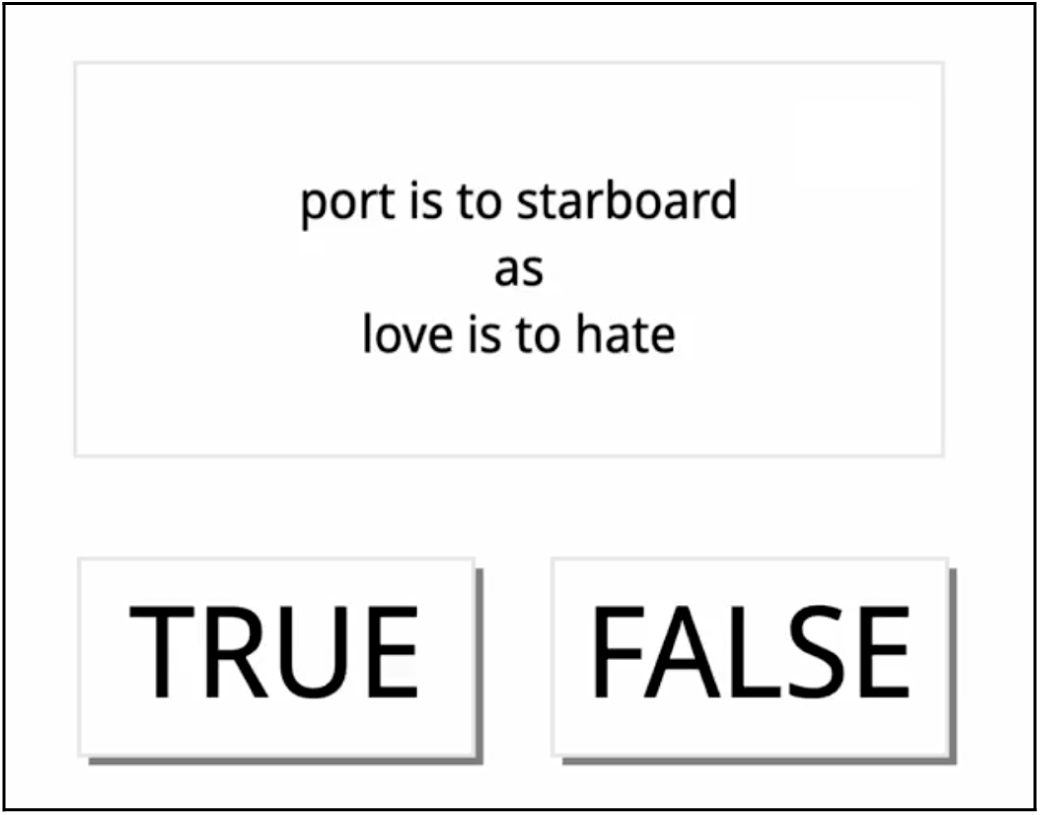
A trial in the Verbal Analogies task.

**Figure 10.**
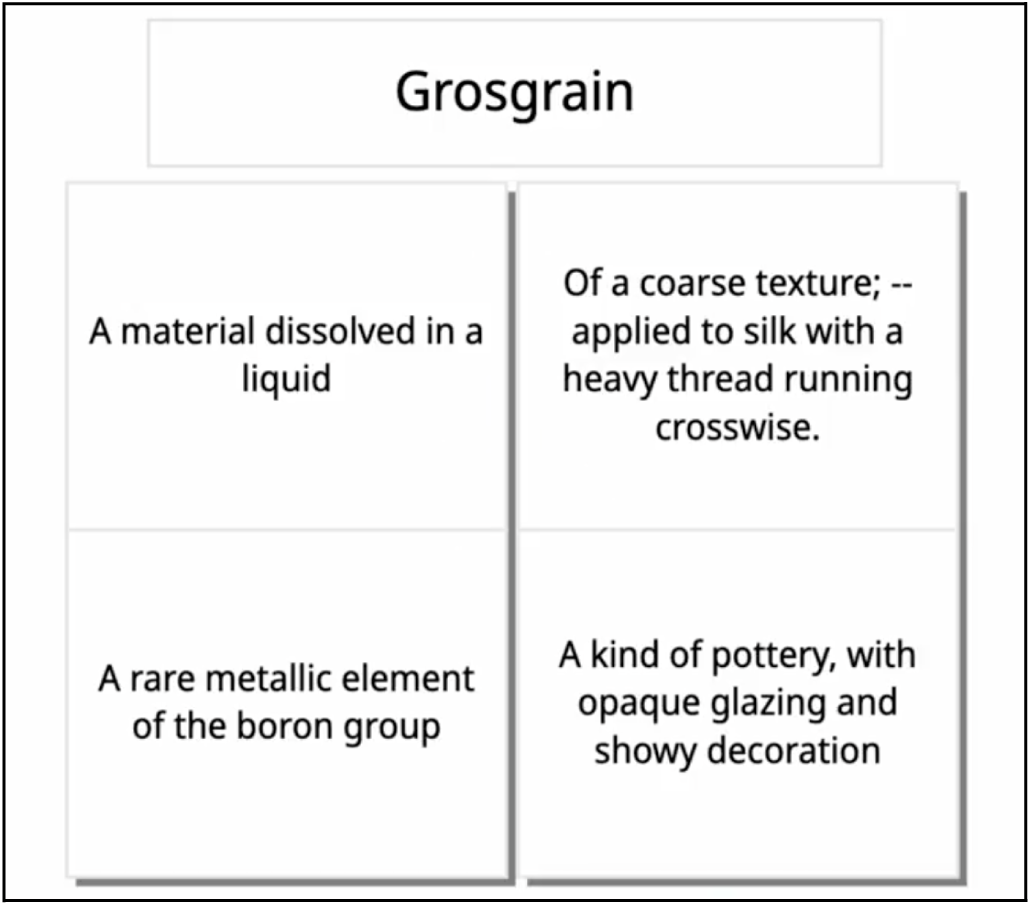
One trial in the Word Definitions task.

**Figure 11.**
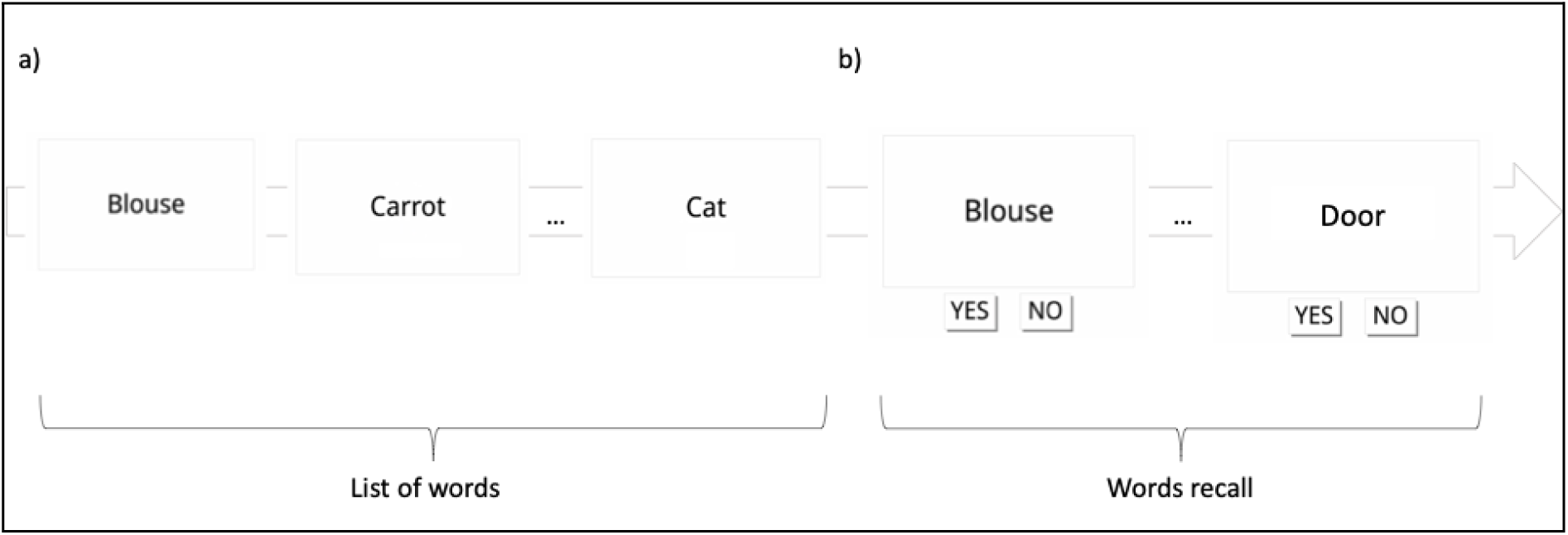
Schematic representation of *Word Memory Immediate and Delayed*. Panel *a)* shows the list of words presented to the participant in the first part of the task. Panel *b)* shows the recall where participants responded whether the word was previously presented in the list or not.

#### The sequence and duration of movements for the somatotopic mapping task

**Table 1.** Sequence of trials for run 1 of the somatotopy task.

| Movement | Duration (seconds) |
| --- | --- |
| right hand | 18 |
| right foot | 17 |
| left tongue | 15 |
| left hand | 15 |
| left foot | 18 |
| right part of the face | 17 |
| rest | 20 |
| right tongue | 18 |
| left hand | 18 |
| right part of the face | 15 |
| right foot | 18 |
| right tongue | 17 |
| left part of the face | 17 |
| rest | 18 |
| left hand | 17 |
| right hand | 16 |
| left tongue | 18 |
| right part of the face | 18 |
| right foot | 15 |
| left foot | 15 |
| rest | 22 |
| left tongue | 15 |
| left part of the face | 15 |
| right hand | 16 |
| left part of the face | 18 |
| left foot | 17 |
| right tongue | 16 |

**Table 2.**
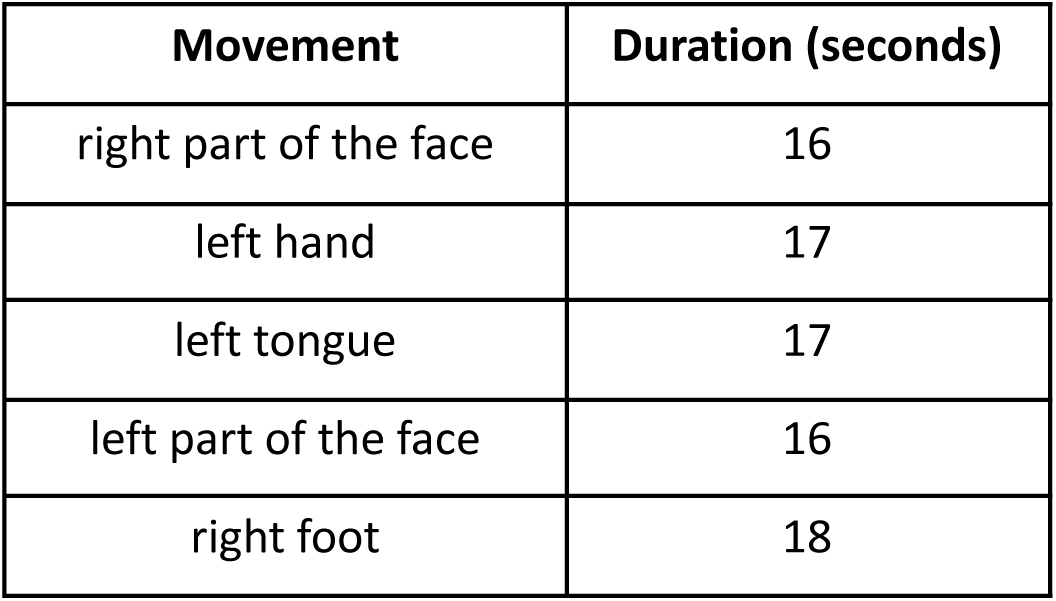
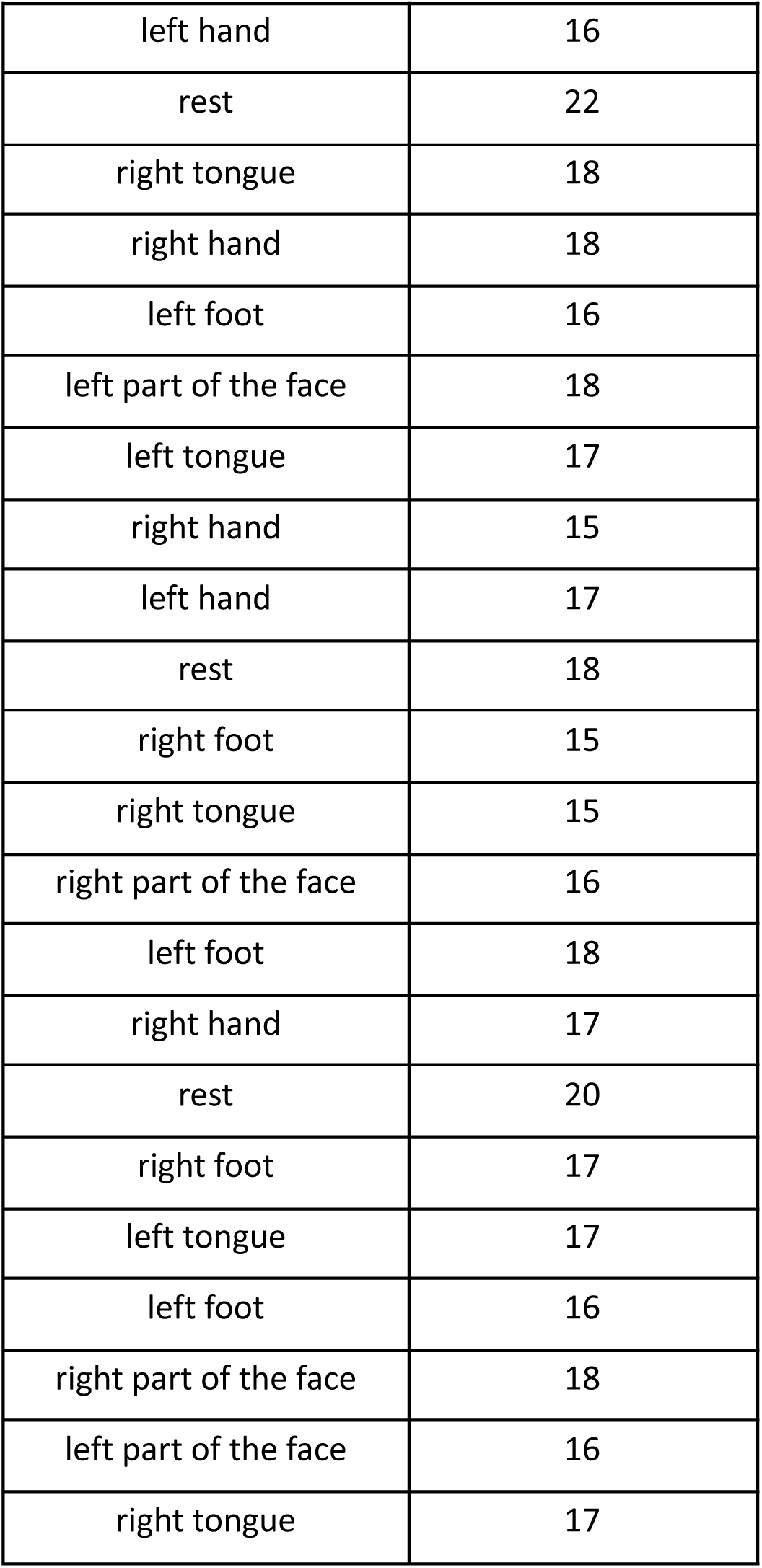
Sequence of trials for run 1 of the somatotopy task.

#### Eye-tracker calibration quality of each participant, eye and run for backtothefuture task

**Table 3.**
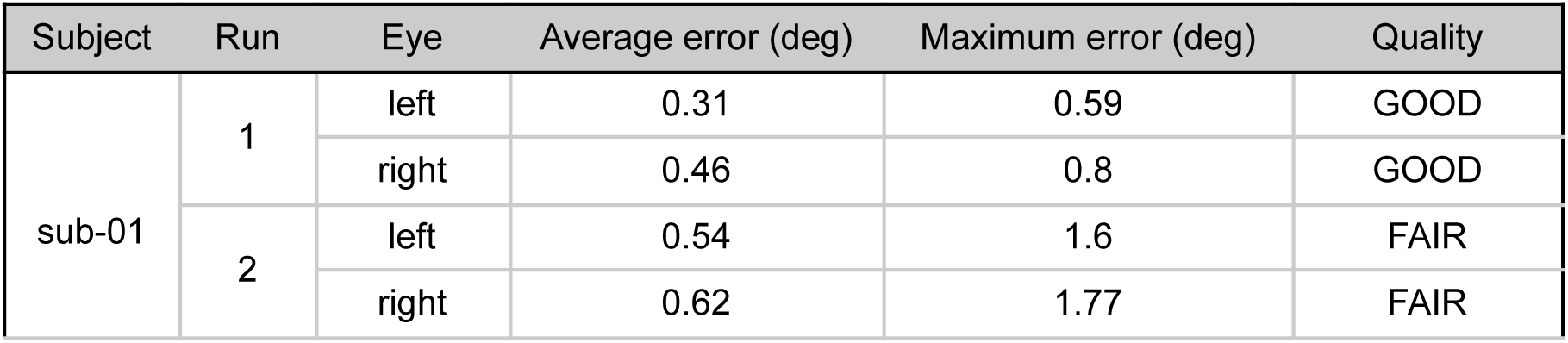
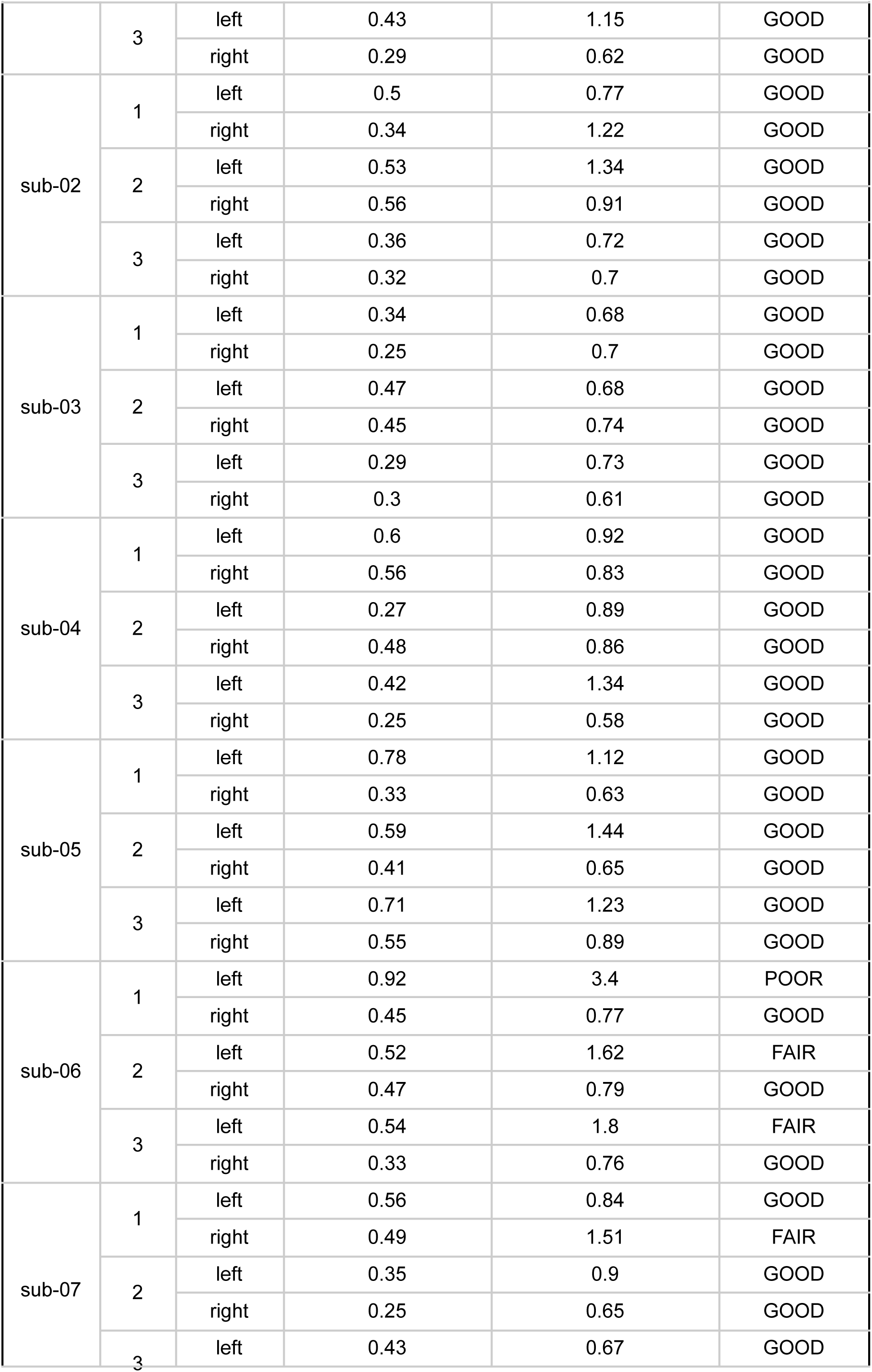
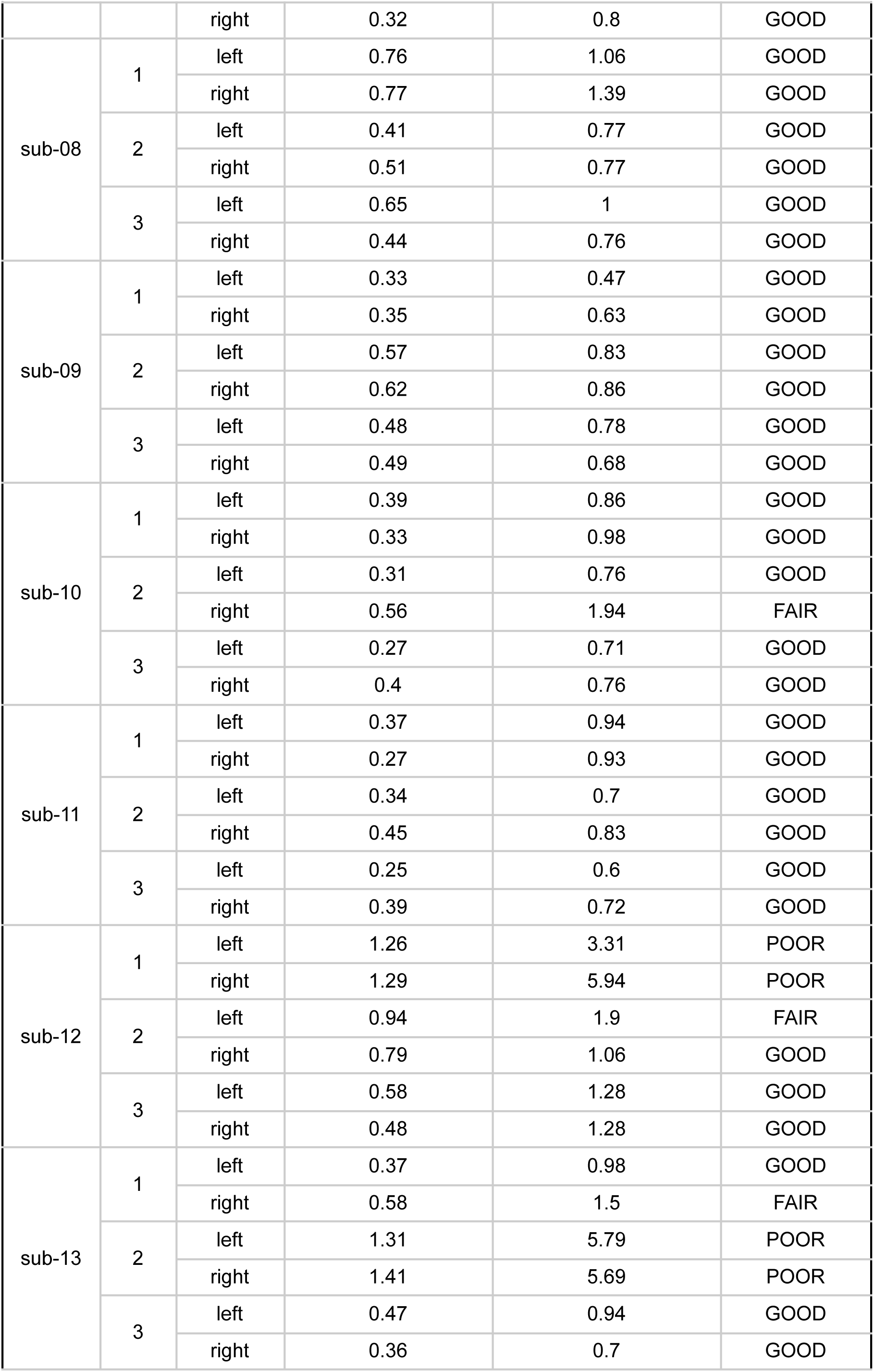
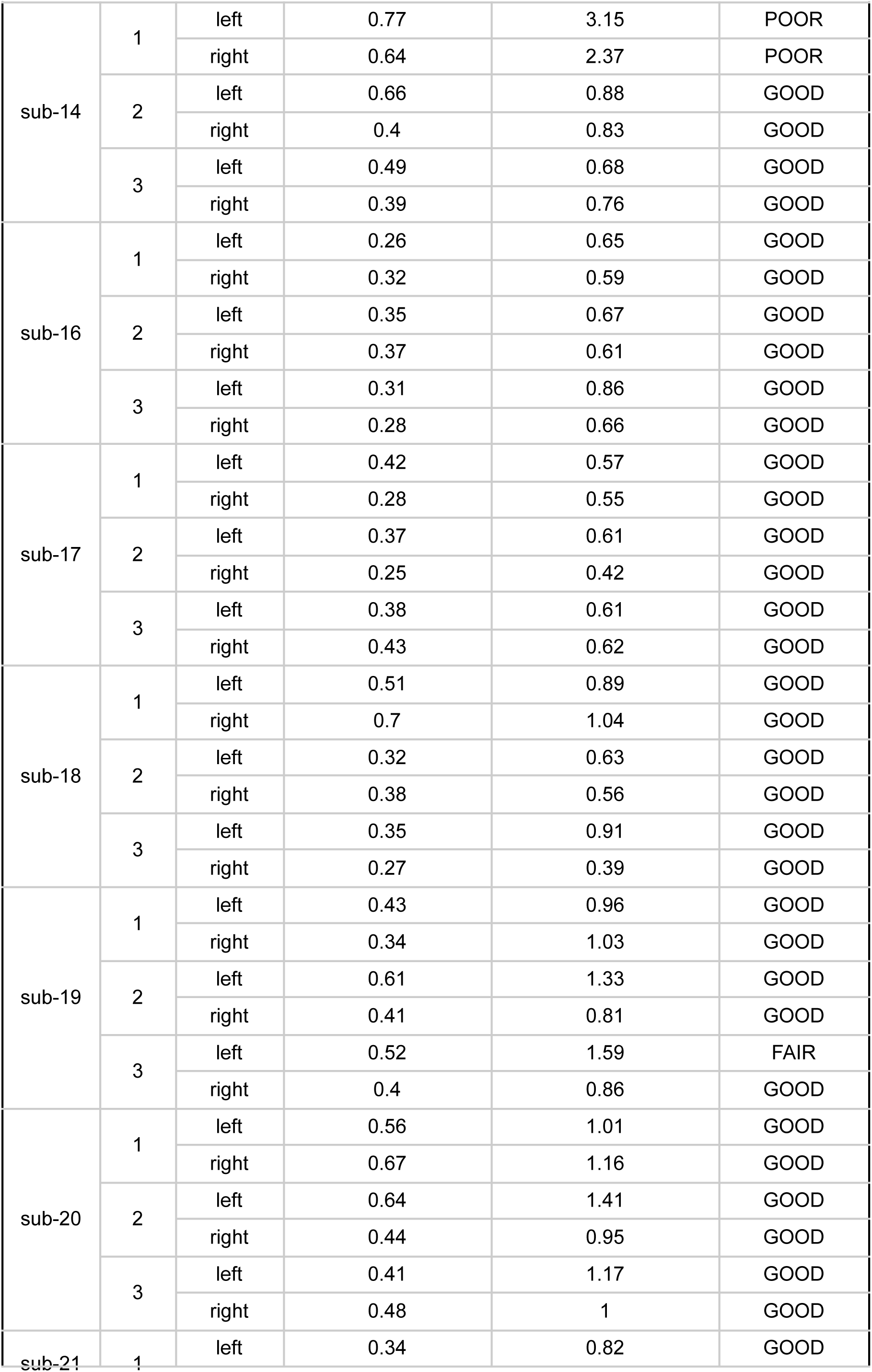
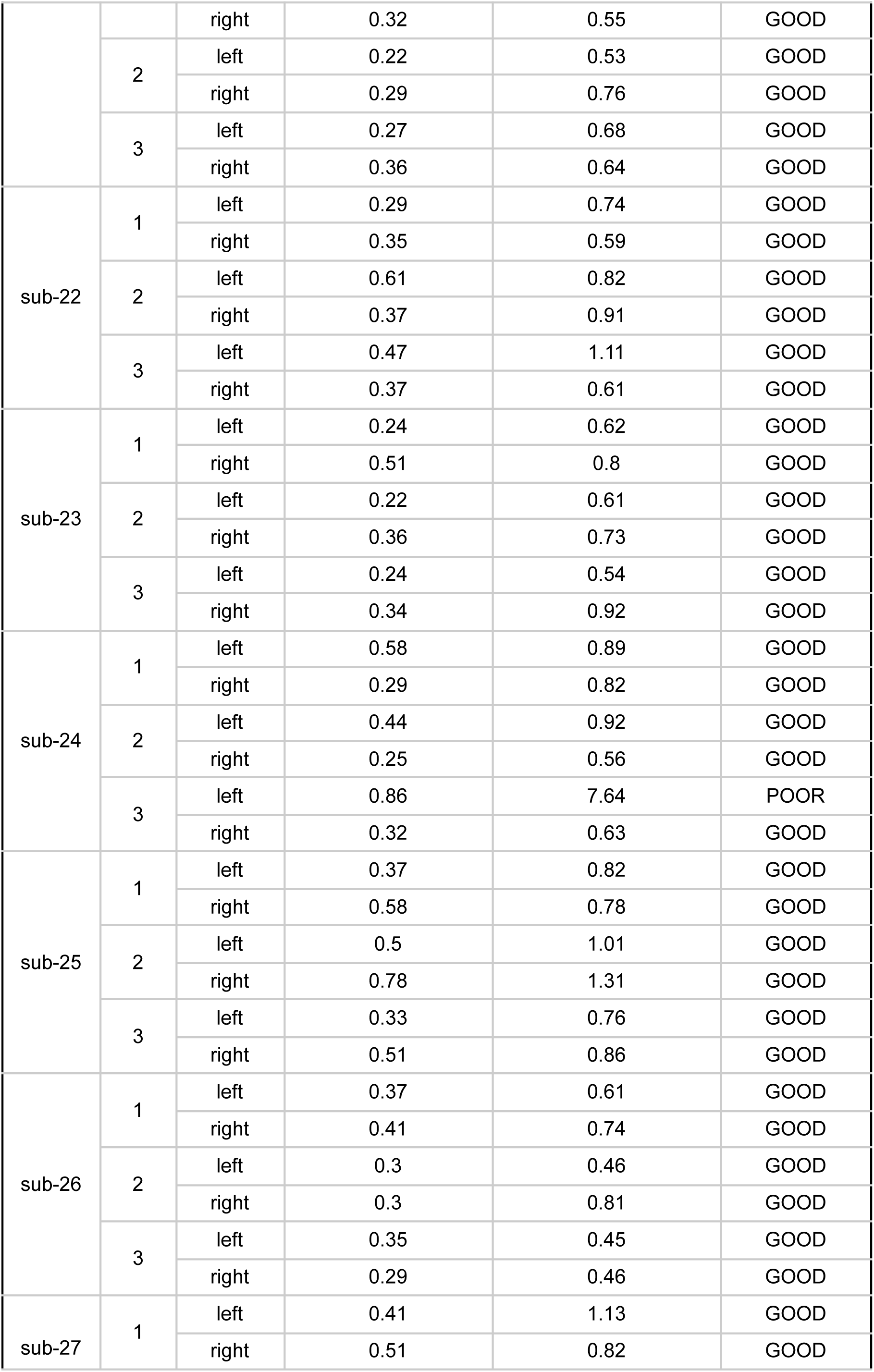
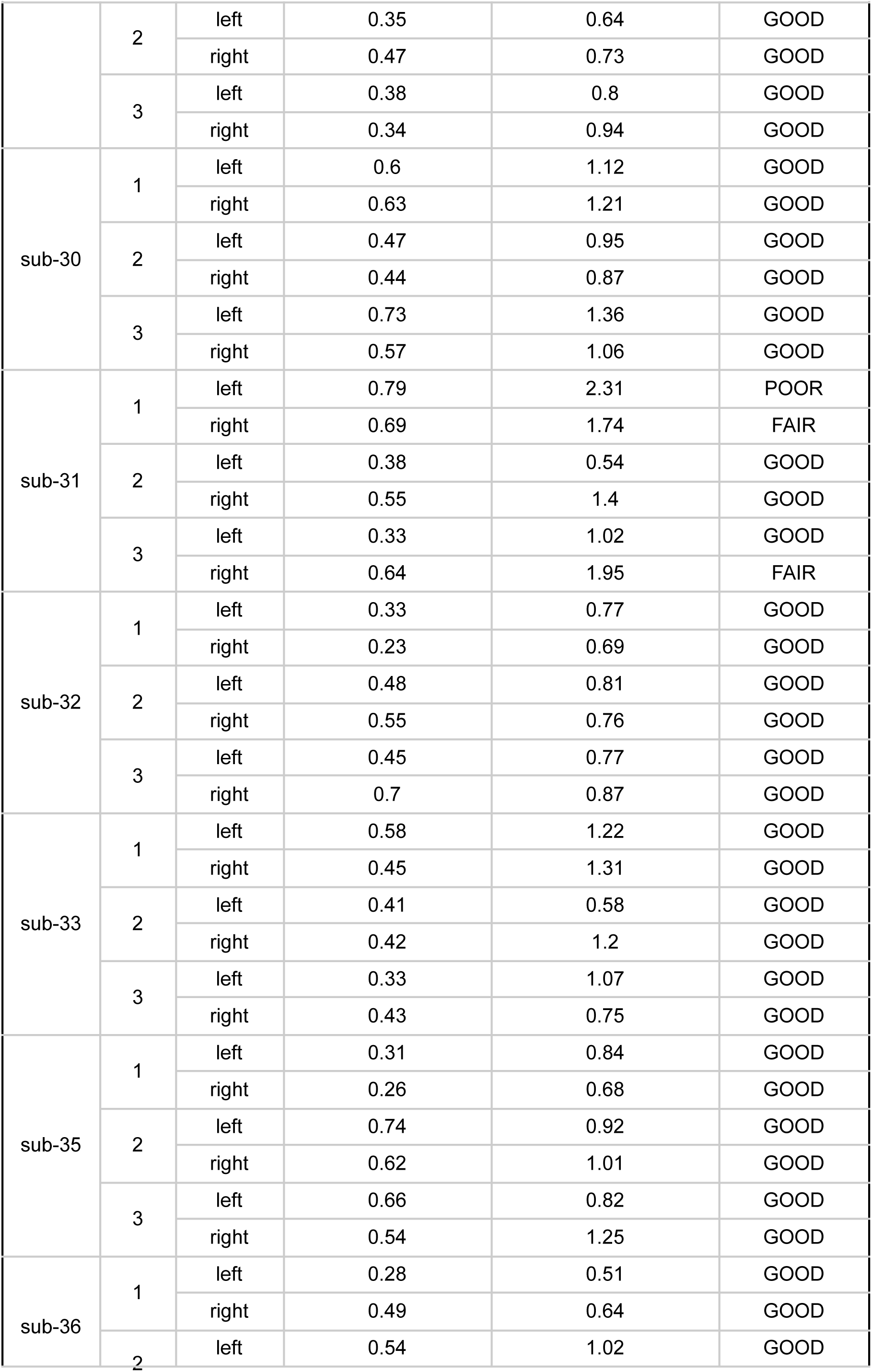
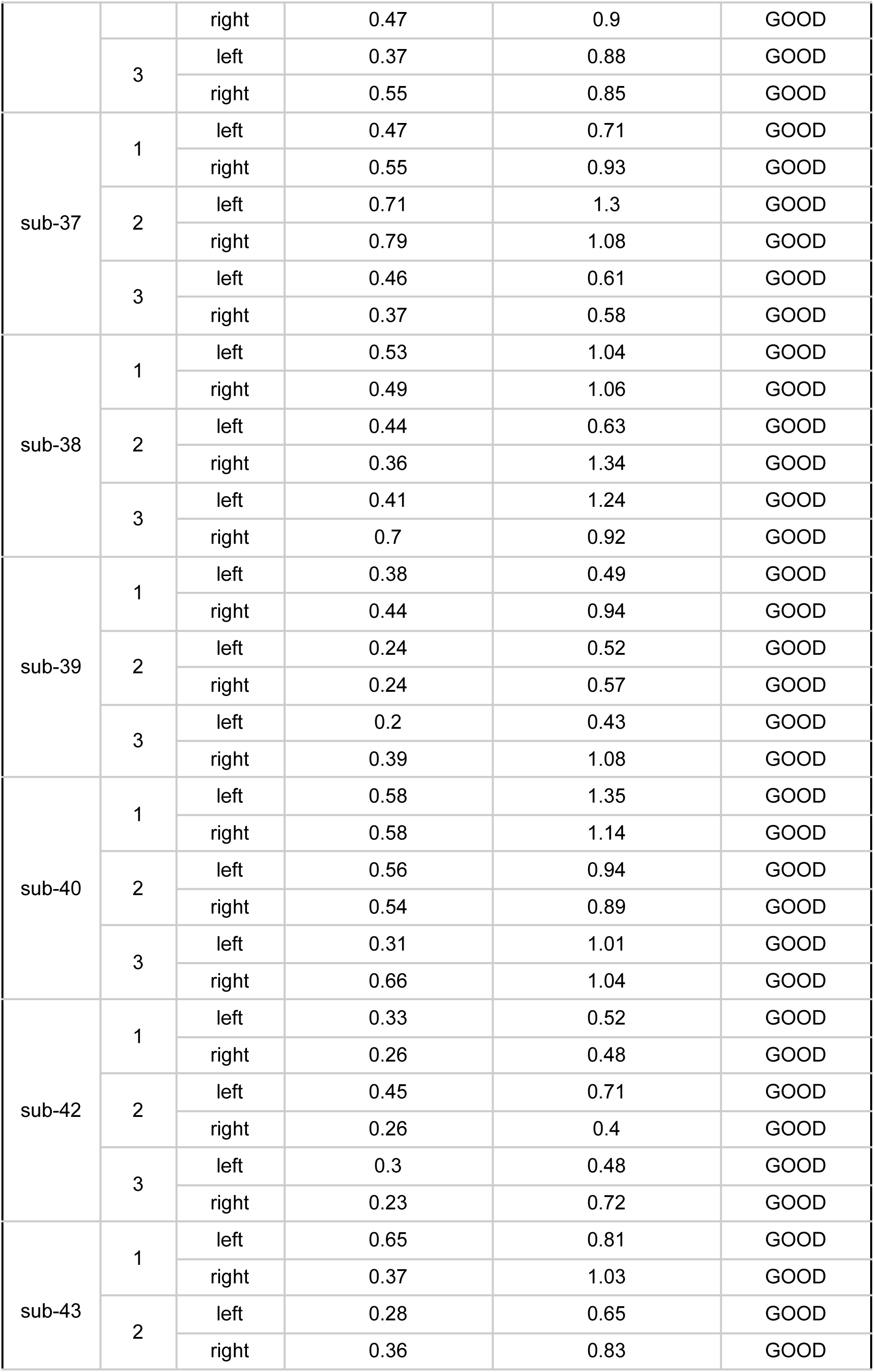
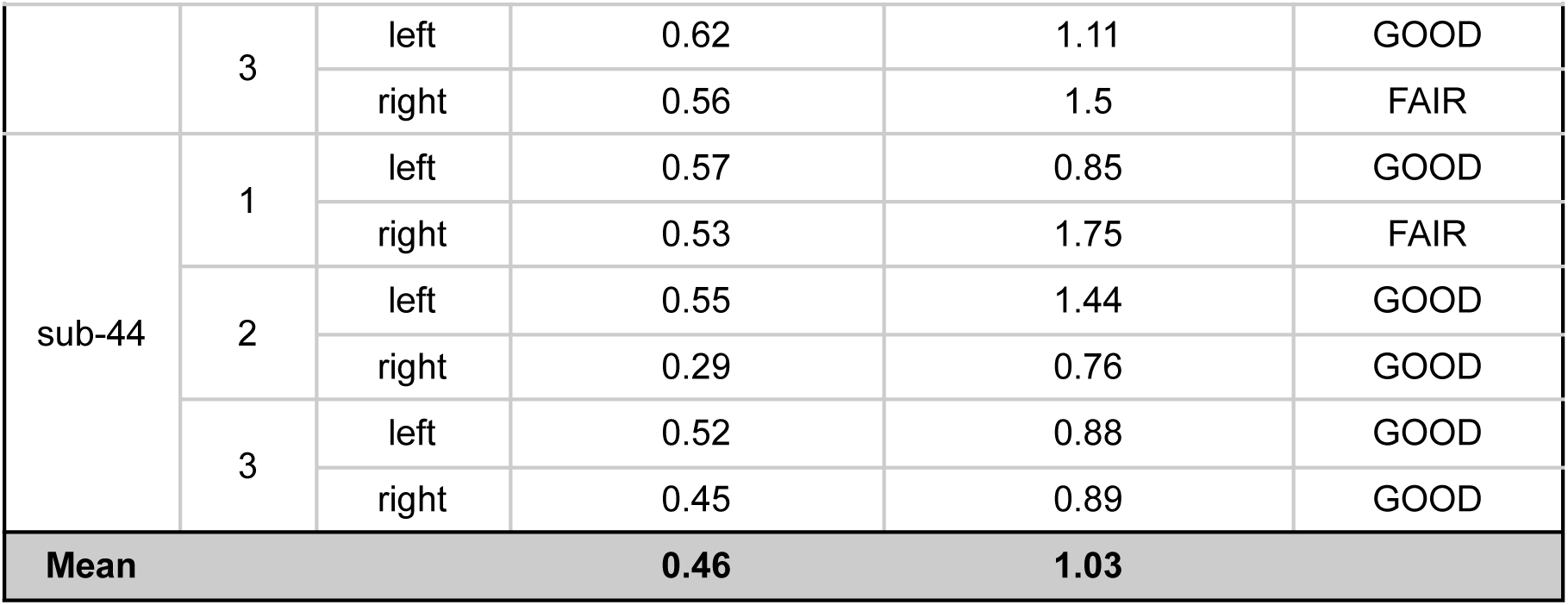
Average and maximum errors in degrees per participant, run, and eye for the backtothefuture task.

## Footnotes

1 https://www.mail-archive.com/freesurfer@nmr.mgh.harvard.edu/msg38136.html

## Notes

### Competing Interest Statement

The authors have declared no competing interest.

### Summary of Updates

Figure 1 revised, typos corrected, more details on cognitive tasks provided

## References

Alderson-Day, B., Mitrenga, K., Wilkinson, S., McCarthy-Jones, S., & Fernyhough, C. (2018). The varieties of inner speech questionnaire – Revised (VISQ-R): Replicating and refining links between inner speech and psychopathology. Consciousness and Cognition, 65, 48–58. 10.1016/j.concog.2018.07.001

Aliko, S., Huang, J., Gheorghiu, F., Meliss, S., & Skipper, J. I. (2020). A naturalistic neuroimaging database for understanding the brain using ecological stimuli. Scientific Data, 7(1), 347. 10.1038/s41597-020-00680-2

Baer, R. A., Smith, G. T., Lykins, E., Button, D., Krietemeyer, J., Sauer, S., Walsh, E., Duggan, D., & Williams, J. M. G. (2008). Construct Validity of the Five Facet Mindfulness Questionnaire in Meditating and Nonmeditating Samples. Assessment, 15(3), 329–342. 10.1177/1073191107313003

Baldassano, C., Hasson, U., & Norman, K. A. (2018). Representation of Real-World Event Schemas during Narrative Perception. The Journal of Neuroscience, 38(45), 9689–9699. 10.1523/JNEUROSCI.0251-18.2018

Brinthaupt, T. M., Hein, M. B., & Kramer, T. E. (2009). The Self-Talk Scale: Development, Factor Analysis, and Validation. Journal of Personality Assessment, 91(1), 82–92. 10.1080/00223890802484498

Chen, J., Leong, Y. C., Honey, C. J., Yong, C. H., Norman, K. A., & Hasson, U. (2017). Shared memories reveal shared structure in neural activity across individuals. Nature Neuroscience, 20(1), 115–125. 10.1038/nn.4450

Chow-Wing-Bom, H. T., Lisi, M., Benson, N. C., Lygo-Frett, F., Yu-Wai-Man, P., Dick, F., Maimon-Mor, R. O., & Dekker, T. M. (2025). Mapping Visual Contrast Sensitivity and Vision Loss Across the Visual Field with Model-Based fMRI. 10.7554/eLife.105930.1

Cox, R. W. (1996). AFNI: Software for Analysis and Visualization of Functional Magnetic Resonance Neuroimages. Computers and Biomedical Research, 29(3), 162–173. 10.1006/cbmr.1996.0014

Cox, R. W., & Hyde, J. S. (1997). Software tools for analysis and visualization of fMRI data. NMR in Biomedicine, 10(4–5), 171–178. 10.1002/(SICI)1099-1492(199706/08)10:4/5<171::AID-NBM453>3.0.CO;2-L

Dekker, T. M., Schwarzkopf, D. S., De Haas, B., Nardini, M., & Sereno, M. I. (2019). Population receptive field tuning properties of visual cortex during childhood. Developmental Cognitive Neuroscience, 37, 100614. 10.1016/j.dcn.2019.01.001

Del Giovane, M., Trender, W. R., Bălăeţ, M., Mallas, E.-J., Jolly, A. E., Bourke, N. J., Zimmermann, K., Graham, N. S. N., Lai, H., Losty, E. J. F., Oiarbide, G. A., Hellyer, P. J., Faiman, I., Daniels, S. J. C., Batey, P., Harrison, M., Giunchiglia, V., Kolanko, M. A., David, M. C. B., … Hampshire, A. (2023). Computerised cognitive assessment in patients with traumatic brain injury: An observational study of feasibility and sensitivity relative to established clinical scales. eClinicalMedicine, 59, 101980. 10.1016/j.eclinm.2023.101980

Destrieux, C., Fischl, B., Dale, A., & Halgren, E. (2010). Automatic parcellation of human cortical gyri and sulci using standard anatomical nomenclature. NeuroImage, 53(1), 1–15. 10.1016/j.neuroimage.2010.06.010

Dick, F. K., Lehet, M. I., Callaghan, M. F., Keller, T. A., Sereno, M. I., & Holt, L. L. (2017). Extensive Tonotopic Mapping across Auditory Cortex Is Recapitulated by Spectrally Directed Attention and Systematically Related to Cortical Myeloarchitecture. The Journal of Neuroscience, 37(50), 12187–12201. 10.1523/JNEUROSCI.1436-17.2017

Dick, F., Taylor Tierney, A., Lutti, A., Josephs, O., Sereno, M. I., & Weiskopf, N. (2012). *In Vivo* Functional and Myeloarchitectonic Mapping of Human Primary Auditory Areas. The Journal of Neuroscience, 32(46), 16095–16105. 10.1523/JNEUROSCI.1712-12.2012

Fischl, B. (2012). FreeSurfer. NeuroImage, 62(2), 774–781. 10.1016/j.neuroimage.2012.01.021

Gal, S., Coldham, Y., Tik, N., Bernstein-Eliav, M., & Tavor, I. (2022). Act natural: Functional connectivity from naturalistic stimuli fMRI outperforms resting-state in predicting brain activity. NeuroImage, 258, 119359. 10.1016/j.neuroimage.2022.119359

Gorgolewski, K. J., Auer, T., Calhoun, V. D., Craddock, R. C., Das, S., Duff, E. P., Flandin, G., Ghosh, S. S., Glatard, T., Halchenko, Y. O., Handwerker, D. A., Hanke, M., Keator, D., Li, X., Michael, Z., Maumet, C., Nichols, B. N., Nichols, T. E., Pellman, J., … Poldrack, R. A. (2016). The brain imaging data structure, a format for organizing and describing outputs of neuroimaging experiments. Scientific Data, 3(1), 160044. 10.1038/sdata.2016.44

Grant, D. A., & Berg, E. (1948). A behavioral analysis of degree of reinforcement and ease of shifting to new responses in a Weigl-type card-sorting problem. Journal of Experimental Psychology, 38(4), 404–411. 10.1037/h0059831

Hanke, M., Baumgartner, F. J., Ibe, P., Kaule, F. R., Pollmann, S., Speck, O., Zinke, W., & Stadler, J. (2014). A high-resolution 7-Tesla fMRI dataset from complex natural stimulation with an audio movie. Scientific Data, 1(1), 140003. 10.1038/sdata.2014.3

Hasson, U., Nir, Y., Levy, I., Fuhrmann, G., & Malach, R. (2004). Intersubject Synchronization of Cortical Activity During Natural Vision. Science, 303(5664), 1634–1640. 10.1126/science.1089506

Heavey, C. L., Moynihan, S. A., Brouwers, V. P., Lapping-Carr, L., Krumm, A. E., Kelsey, J. M., Turner, D. K., & Hurlburt, R. T. (2019). Measuring the Frequency of Inner-Experience Characteristics by Self-Report: The Nevada Inner Experience Questionnaire. Frontiers in Psychology, 9, 2615. 10.3389/fpsyg.2018.02615

Hutchison, R. M., Womelsdorf, T., Allen, E. A., Bandettini, P. A., Calhoun, V. D., Corbetta, M., Della Penna, S., Duyn, J. H., Glover, G. H., Gonzalez-Castillo, J., Handwerker, D. A., Keilholz, S., Kiviniemi, V., Leopold, D. A., de Pasquale, F., Sporns, O., Walter, M., & Chang, C. (2013). Dynamic functional connectivity: Promise, issues, and interpretations. NeuroImage, 80, 360–378. 10.1016/j.neuroimage.2013.05.079

Ki, J. J., Kelly, S. P., & Parra, L. C. (2016). Attention Strongly Modulates Reliability of Neural Responses to Naturalistic Narrative Stimuli. The Journal of Neuroscience, 36(10), 3092–3101. 10.1523/JNEUROSCI.2942-15.2016

Kroenke, K., Spitzer, R. L., & Williams, J. B. W. (2001). The PHQ-9: Validity of a brief depression severity measure. Journal of General Internal Medicine, 16(9), 606–613. 10.1046/j.1525-1497.2001.016009606.x

Lerner, Y., Honey, C. J., Silbert, L. J., & Hasson, U. (2011). Topographic Mapping of a Hierarchy of Temporal Receptive Windows Using a Narrated Story. The Journal of Neuroscience, 31(8), 2906–2915. 10.1523/JNEUROSCI.3684-10.2011

Mehling, W. E., Acree, M., Stewart, A., Silas, J., & Jones, A. (2018). The Multidimensional Assessment of Interoceptive Awareness, Version 2 (MAIA-2). PLOS ONE, 13(12), e0208034. 10.1371/journal.pone.0208034

Nastase, S. A., Gazzola, V., Hasson, U., & Keysers, C. (2019). Measuring shared responses across subjects using intersubject correlation. Social Cognitive and Affective Neuroscience, 14(6), 667–685. 10.1093/scan/nsz037

Nastase, S. A., Liu, Y.-F., Hillman, H., Norman, K. A., & Hasson, U. (2020). Leveraging shared connectivity to aggregate heterogeneous datasets into a common response space. NeuroImage, 217, 116865. 10.1016/j.neuroimage.2020.116865

Nastase, S. A., Liu, Y.-F., Hillman, H., Zadbood, A., Hasenfratz, L., Keshavarzian, N., Chen, J., Honey, C. J., Yeshurun, Y., Regev, M., Nguyen, M., Chang, C. H. C., Baldassano, C., Lositsky, O., Simony, E., Chow, M. A., Leong, Y. C., Brooks, P. P., Micciche, E., … Hasson, U. (2021). The “Narratives” fMRI dataset for evaluating models of naturalistic language comprehension. Scientific Data, 8(1), 250. 10.1038/s41597-021-01033-3

Nguyen, V. T., Sonkusare, S., Stadler, J., Hu, X., Breakspear, M., & Guo, C. C. (2017). Distinct Cerebellar Contributions to Cognitive-Perceptual Dynamics During Natural Viewing. Cerebral Cortex, 27(12), 5652–5662. 10.1093/cercor/bhw334

Redcay, E., & Moraczewski, D. (2020). Social cognition in context: A naturalistic imaging approach. NeuroImage, 216, 116392. 10.1016/j.neuroimage.2019.116392

Ren, Y., Nguyen, V. T., Guo, L., & Guo, C. C. (2017). Inter-subject Functional Correlation Reveal a Hierarchical Organization of Extrinsic and Intrinsic Systems in the Brain. Scientific Reports, 7(1), 10876. 10.1038/s41598-017-11324-8

Sereno, M. I., Dale, A. M., Reppas, J. B., Kwong, K. K., Belliveau, J. W., Brady, T. J., Rosen, B. R., & Tootell, R. B. H. (1995). Borders of Multiple Visual Areas in Humans Revealed by Functional Magnetic Resonance Imaging. Science, 268(5212), 889–893. 10.1126/science.7754376

Shafto, M. A., Tyler, L. K., Dixon, M., Taylor, J. R., Rowe, J. B., Cusack, R., Calder, A. J., Marslen-Wilson, W. D., Duncan, J., Dalgleish, T., Henson, R. N., Brayne, C., Matthews, F. E., & Cam-CAN. (2014). The Cambridge Centre for Ageing and Neuroscience (Cam-CAN) study protocol: A cross-sectional, lifespan, multidisciplinary examination of healthy cognitive ageing. BMC Neurology, 14, 204. 10.1186/s12883-014-0204-1

Simony, E., Honey, C. J., Chen, J., Lositsky, O., Yeshurun, Y., Wiesel, A., & Hasson, U. (2016). Dynamic reconfiguration of the default mode network during narrative comprehension. Nature Communications, 7(1), 12141. 10.1038/ncomms12141

Spitzer, R. L., Kroenke, K., Williams, J. B. W., & Löwe, B. (2006). A Brief Measure for Assessing Generalized Anxiety Disorder: The GAD-7. Archives of Internal Medicine, 166(10), 1092. 10.1001/archinte.166.10.1092

Stroop, J. R. (1935). Studies of interference in serial verbal reactions. Journal of Experimental Psychology, 18(6), 643–662. 10.1037/h0054651

Tennant, R., Hiller, L., Fishwick, R., Platt, S., Joseph, S., Weich, S., Parkinson, J., Secker, J., & Stewart-Brown, S. (2007). The Warwick-Edinburgh Mental Well-being Scale (WEMWBS): Development and UK validation. Health and Quality of Life Outcomes, 5(1), 63. 10.1186/1477-7525-5-63

Vanderwal, T., Eilbott, J., & Castellanos, F. X. (2019). Movies in the magnet: Naturalistic paradigms in developmental functional neuroimaging. Developmental Cognitive Neuroscience, 36, 100600. 10.1016/j.dcn.2018.10.004

Visconti Di Oleggio Castello, M., Chauhan, V., Jiahui, G., & Gobbini, M. I. (2020). An fMRI dataset in response to “The Grand Budapest Hotel”, a socially-rich, naturalistic movie. Scientific Data, 7(1), 383. 10.1038/s41597-020-00735-4

Wegner, D. M., & Zanakos, S. (1994). Chronic Thought Suppression. Journal of Personality, 62(4), 615–640. 10.1111/j.1467-6494.1994.tb00311.x

Xu, J., Moeller, S., Auerbach, E. J., Strupp, J., Smith, S. M., Feinberg, D. A., Yacoub, E., & Uğurbil, K. (2013). Evaluation of slice accelerations using multiband echo planar imaging at 3T. NeuroImage, 83, 991–1001. 10.1016/j.neuroimage.2013.07.055

Zemeckis, R. (Director). (1985). Back to the Future [Video recording].

